# Global Huntingtin Knockout in Adult Mice Leads to Fatal Neurodegeneration that Spares the Pancreas

**DOI:** 10.1101/2024.01.11.575238

**Authors:** Robert M. Bragg, Ella W. Mathews, Andrea Grindeland, Jeffrey P. Cantle, David Howland, Tom Vogt, Jeffrey B. Carroll

## Abstract

Huntington’s disease (HD) is a fatal neurogenerative disorder caused by an expanded glutamine-coding CAG tract in the Huntingtin (Htt) gene. HD is believed to primarily arise via a toxic gain of function, and as a result a wide range of Htt-lowering treatments are in clinical trials. The safety of these trials is contingent on the risks imposed by Htt lowering: Htt is widely conserved, ubiquitously expressed and its complete loss causes severe developmental symptoms in mice and humans. Recently, multiple labs have reported on the consequences of widespread inducible Htt loss in mice. One report describes that early induction of global Htt loss causes fatal pancreatitis, but that later onset lowering is benign. Another study did not report fatal pancreatitis but suggested that postnatal Htt loss was associated with widespread progressive phenotypes, including subcortical calcification and neurodegeneration. To better understand the risks posed by widespread inducible Htt loss we established the phenotypes of mice in which we knocked out Htt with two tamoxifen inducible Cre lines, which we have here extensively characterized. In short, we find that widespread loss of Htt at 2 months of age leads to a wide range of phenotypes, including subcortical calcification, but does not result in acute pancreatitis or histological changes in the pancreas. Additionally, we report here for the first time that Htt loss is followed by robust and sustained increases in the levels of neurofilament light chain (NfL), a peripherally accessible biomarker of neuroaxonal stress. These results confirm that complete loss of Htt in mice is associated with pronounced risks, including progressive subcortical calcification and neurodegeneration.

## Introduction

The Huntingtin (*Htt*) gene is widely conserved and ubiquitously expressed across investigated cell types. *Htt* is most well known as the gene in which an expansion of a glutamine-coding CAG repeat causes Huntington’s disease (HD), an autosomal dominant, fatal, neurodegenerative disease (1). *Htt* expression appears to be very important, particularly during development, as *Htt* null mice die very early in embryogenesis (2–4), and compound heterozygous expression of hypomorphic *Htt* alleles causes a profound neurodevelopmental impairment in humans, with non-progressive features distinct from HD (5,6). Population databases reveal that putative loss of function (pLoF) mutations in *Htt* are highly selected against (e.g. the abundance of observed/expected pLoF mutations in gnomAD (7) is = 0.12; pLI score = 1).

Genetic evidence suggests that HD pathology primarily arises from a toxic gain of function (1). This, coupled with dominant and complete penetrance, makes Huntingtin (HTT)-lowering therapies an attractive approach to HD treatment (8). Indeed, a wide range of HTT-lowering approaches have already advanced to human clinical studies, including ongoing studies with antisense oligonucleotides (ASOs), virally delivered microRNA (miRNAs), and most recently small molecule splice modulators that favor the inclusion of a cryptic exon with a premature stop codon, thereby reducing HTT levels across the body (9,10). Some of these approaches are inherently allele-selective, only targeting mutant HTT (mHTT), for example intrathecally delivered ASOs developed by Wave Life Sciences which target single nucleotide polymorphisms (SNPs) in the *Htt* gene (11). Other approaches, for example Tominersen – an ASO targeting both wildtype and mutant *Htt*, currently being developed by Ionis Pharmaceuticals and Roche (11,12) - target both alleles. Genetic evidence suggests that 50% HTT expression is well tolerated in both mice and humans - the heterozygous parents of patients impacted by a syndrome caused by expression of very low levels of *HTT* are not described to have any clinically meaningful impact of their near 50% HTT expression. Similarly, in mice, while complete knockout (KO) of *Htt* results in embryonic lethality, heterozygous KO mice are grossly normal – to our knowledge, the only reported phenotype in these mice is a reduction in body weight (13). If HTT-lowering therapies work, a key question about their safety is the extent to which HTT silencing is tolerated, and the precise level of HTT required for healthy cellular function.

A key piece of information needed to assess the safety of HTT lowering is the tolerability of wide-spread, adult-onset HTT-lowering. Indeed, two groups have published studies with similar designs that investigated the impact of tamoxifen-Cre-induced recombination and global HTT lowering in *Htt-*flox mice (14,15). Of these, Dietrich et al. (2017), found widespread deleterious effects of adult onset HTT lowering, including progressive neurodegeneration accompanied by thalamic calcification (15). In contrast, Wang et al. (2016) found that global HTT silencing at 4 and 8 months was benign through 18 months of age, but that silencing HTT at 2 months of age led to acute and fatal pancreatitis, with 95% fatality by 10 days of *Htt* loss (14). Motivated to better understand these apparently discordant results, we conducted a replication study.

We designed our study to closely mimic the two previous ones, with several additional assessments. We generated two separate cohorts of *Htt-*flox mice crossed with unique ubiquitous tamoxifen inducible Cre recombinase (Cre) lines. In the floxed *Htt* allele, Cre expression deletes the entire first exon, including the transcription start site, as well as the promoter and part of intron 1. We first evaluated the leakiness – the tendency to recombine the targeted locus in the absence of tamoxifen treatment – in both Cre lines across a wide range of tissues using quantitative *Htt* mRNA and protein assays. We then initiated tamoxifen treatment to remove HTT expression at beginning at 2 months of age and lasting until sacrifice at 14 months of age. In contrast to Wang et al., we did not observe pancreatis after early-onset HTT loss. In addition to behavioral and pathological endpoints following extended HTT silencing, we also collected longitudinal plasma samples to measure levels of neurofilament light chain (NfL) - a widely used biomarker for neuronal stress (16). Strikingly, we observed progressive neurological changes including the appearance of calcified deposits in the thalamus, confirming findings reported by Dietrich et al., and discovered that plasma NfL is robustly increased by HTT loss.

In summary, we have conducted a replication study to establish the impact of widespread adult-onset HTT lowering. Our results are largely consistent with those of Dietrich et al (2017), while we observe significant differences from those of Wang et al (2016), notably an absence of acute fatal pancreatitis after early *Htt* deletion. Our results support the hypothesis that complete loss of *Htt* in adult animals is poorly tolerated as it is associated with a range of phenotypes, including progressive neurodegeneration accompanied by thalamic calcification.

## Results

### Establishment of Allele Utility

Both previous studies relied on the B6.Cg-Tg(CAG-cre/Esr1*)5Amc/J, referred to as “CAG-Cre” mice (Jackson labs strain #004682), in which a tamoxifen-inducible Cre construct is under the control of a ubiquitous promoter, leading to widespread recombination of floxed alleles after tamoxifen treatment (17). We also used a second ubiquitously expressed tamoxifen-inducible Cre line, which has been shown to have widespread efficiency, including in the brain (18) (B6.Cg-Ndor1^Tg(UBC-cre/ERT2)1Ejb/1^J), referred to as “UBC-Cre” (19). Both lines were crossed to a previously described floxed *Htt* allele, B6.Htt^tm2Szi^ (4), referred to as *Htt^fl/fl^*, resulting in multiple large cohorts of *Htt^fl/fl^*;CAG-Cre, *Htt^fl/fl^*;UBC-Cre and control lines, *Htt^+/+^* and *Htt^fl/fl^*, on a consistent C57Bl/6J background (Table 1), referred to here as conditional *Htt* knock-out (cKO) mice.

**Table 1.**
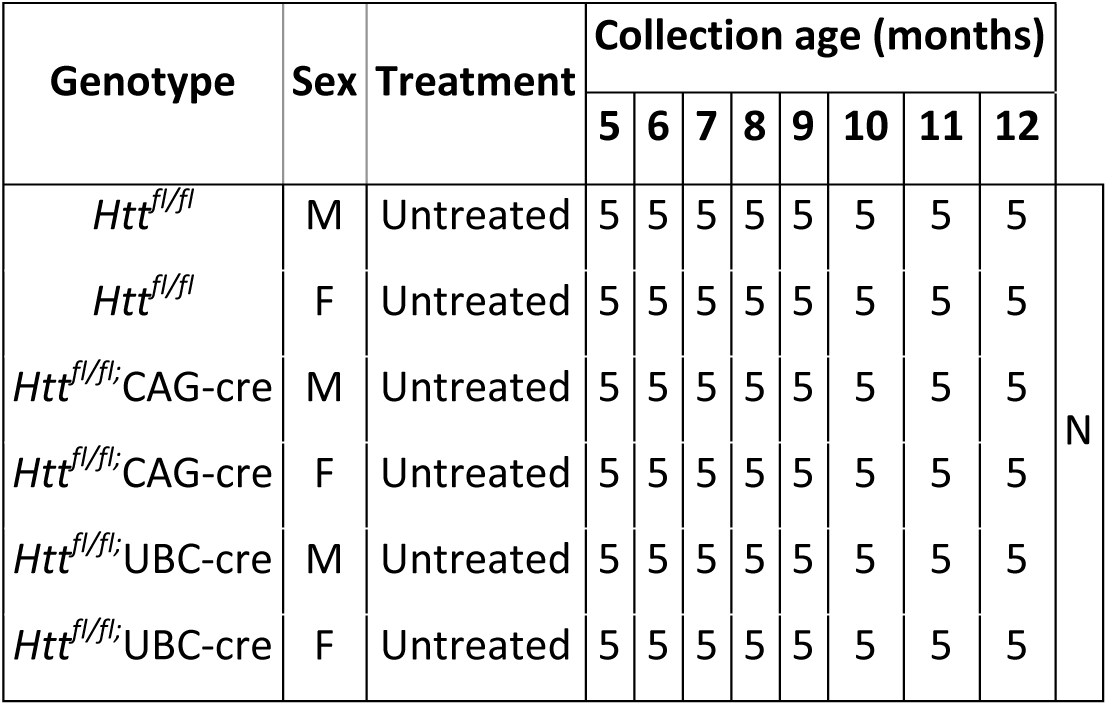
Leak assessment cohort.

We next investigated the leakiness of each Cre construct in the absence of tamoxifen from 5-12 months of age by quantifying the levels of *Htt*, HTT and the estimated percentage recombination frequency using qRT-PCR, electro-chemiluminescent ELISA, and semi-quantitative PCR, respectively (Table 1). In general, our DNA, RNA and protein assays are consistent throughout this study. Interestingly, we observed that the *Htt^fl/fl^*;UBC-Cre line was extremely leaky at the floxed *Htt* locus in the CNS organs examined, with up to 83% reduction in HTT levels in the cortex at 12 months of age (Fig. 1B, Tukey’s post-hoc p < 0.0001 ), but was much more tightly regulated in the examined peripheral organs. In contrast, the *Htt^fl/fl^*;CAG-Cre exhibited more leak in peripheral organs, reaching up to a 53% reduction in HTT in the liver (Fig. 1B, Tukey’s post-hoc p, *p* = 0.002), but was very tightly regulated in CNS tissue. We replicated these observations in a separate follow-up cohort of mice in which we took a wider tissue survey at a single time point (cortex, striatum, cerebellum, spinal cord, liver, kidney, pancreas, and gastrocnemius at 5 months of age; Fig. S1).

**Figure 1.**
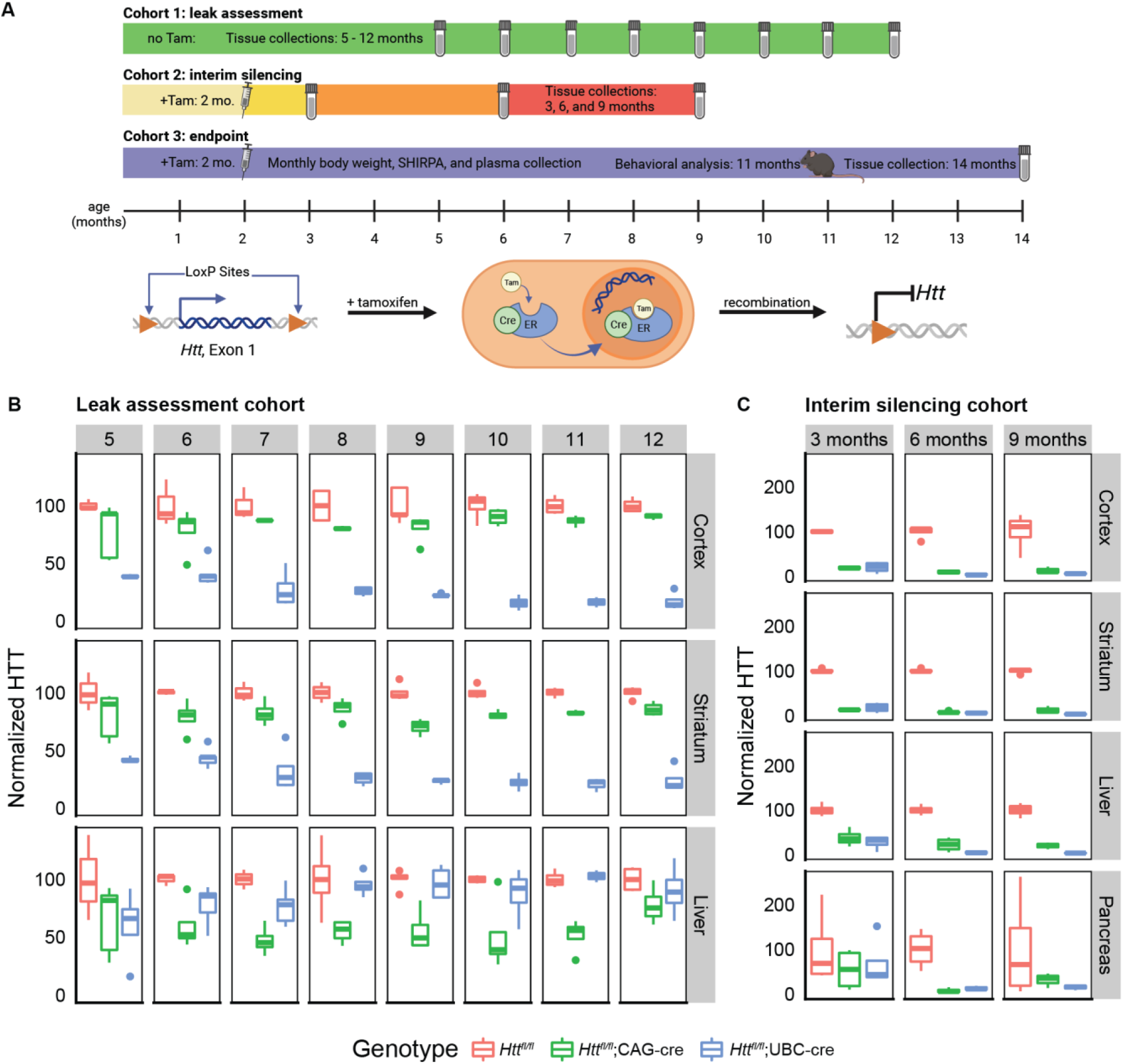
Study design and mouse validation. **A)** (Top) Timelines of experimental cohorts utilized in this study with indications of tamoxifen injection time, in-life analyses, and tissue collection points. (Bottom) Cartoon showing mechanism of Htt inactivation which occurs when tamoxifen (Tam) is administered and causes cytosolic cre-recombinase-estrogen receptor complex to translocate to the nucleus inducing recombination of the floxed Htt allele. **B)** HTT levels measured in the absence of tamoxifen reveal differential leakiness of each Cre line. Column labels indicate age at sacrifice (in months, 5-12). n=5/genotype/age. **C)** Levels of HTT are robustly and consistently reduced in tamoxifen treated mice. Silencing in tamoxifen-treated animals at 3, 6 and 9 months of age (columns) was assessed across a range of organs (rows). n=4/group. HTT levels in B and C were assayed using chemiluminescent ELISA and normalized to control group (Htt^fl/fl^ mice).

### Establishment of Cohorts, Study Design, Survival Analysis

Based on the observations that *Htt^fl/fl^*;CAG-Cre and *Htt^fl/fl^*;UBC-Cre mice are differentially leaky across tissue types, we generated large cohorts of mice with both constructs to follow longitudinally. We chose to deploy the *Htt^fl/fl^*;CAG-Cre and *Htt^fl/fl^*;UBC-Cre mice for experiments looking for phenotypes arising in CNS, and peripheral organs, respectively. The experimental cohort included 160 mice – 10M/10F from each genotype/treatment arm (Table 2), as well as a smaller group of mice for planned interim assessment at 3-, 6-, and 9-months of age to monitor HTT loss across tissues during the study (Table 3). Following best practices established for tamoxifen treatment by Jackson Laboratories (20), we treated mice in our tamoxifen cohorts with 75mg/kg tamoxifen for 5 consecutive days at 2 months of age. The study design outlined in Fig. 1A indicates the planned interventions and their timing. Based on previous findings by Wang et al (2016) (14), we were concerned that our mice would die from acute pancreatitis in response to HTT loss at 2 months of age, however we saw no acute death in any of our mice. As expected, our interim silencing cohorts at 3-, 6-, and 9-months show very robust HTT loss compared to vehicle controls in cortex, striatum, and liver (Fig. 1C, Tukey HSD p < 0.01 for all tissues). In pancreas, HTT is significantly lowered at 6 months compared to vehicle control (Tukey HSD p < 0.001), but not at 3- or 9-months, likely due to high variability in this tissue. By 12 months, both control and cKO mice began to reach humane endpoint criteria due to treatment resistant ulcerative dermatitis, while only cKO mice also displayed severe progressive tremor. We chose to euthanize all remaining mice at 14 months of age to collect a sufficiently sized cohort of age-matched tissue for robust pathological and molecular characterization.

**Table 2.** Experimental cohort.

| Genotype | Sex | Treatment | Initiated N (2 mo) | Collected N (14 mo) |
| --- | --- | --- | --- | --- |
| *Htt*<sup>+/+</sup> | M | Vehicle | 10 | 10 |
| *Htt*<sup>+/+</sup> | F | Vehicle | 10 | 10 |
| *Htt*<sup>fl/fl</sup> | M | Vehicle | 10 | 10 |
| *Htt*<sup>fl/fl</sup> | F | Vehicle | 10 | 10 |
| *Htt*<sup>fl/fl</sup>;CAG-cre | M | Vehicle | 10 | 10 |
| *Htt*<sup>fl/fl</sup>;CAG-cre | F | Vehicle | 10 | 8 |
| *Htt*<sup>fl/fl</sup>;UBC-cre | M | Vehicle | 10 | 10 |
| *Htt*<sup>fl/fl</sup>;UBC-cre | F | Vehicle | 10 | 10 |
| *Htt*<sup>+/+</sup> | M | Tamoxifen | 10 | 10 |
| *Htt*<sup>+/+</sup> | F | Tamoxifen | 10 | 10 |
| *Htt*<sup>fl/fl</sup> | M | Tamoxifen | 10 | 10 |
| *Htt*<sup>fl/fl</sup> | F | Tamoxifen | 10 | 10 |
| *Htt*<sup>fl/fl</sup>;CAG-cre | M | Tamoxifen | 10 | 9 |
| *Htt*<sup>fl/fl</sup>;CAG-cre | F | Tamoxifen | 10 | 5 |
| *Htt*<sup>fl/fl</sup>;UBC-cre | M | Tamoxifen | 10 | 10 |
| *Htt*<sup>fl/fl</sup>;UBC-cre | F | Tamoxifen | 10 | 10 |

**Table 3.** Interim takedown cohort.

| Genotype | Sex | Treatment | Collection age (months) |  |  |  |
| --- | --- | --- | --- | --- | --- | --- |
|  |  |  | 3 | 6 | 9 |  |
| *Htt<sup>fl/fl</sup>* | M | Tamoxifen | 5 | 5 | 5 | N |
| *Htt<sup>fl/fl</sup>* | F | Tamoxifen | 5 | 5 | 5 |  |
| *Htt<sup>fl/fl</sup>;CAG-cre* | M | Tamoxifen | 5 | 5 | 5 |  |
| *Htt<sup>fl/fl</sup>;CAG-cre* | F | Tamoxifen | 5 | 5 | 5 |  |
| *Htt<sup>fl/fl</sup>;UBC-cre* | M | Tamoxifen | 5 | 5 | 5 |  |
| *Htt<sup>fl/fl</sup>;UBC-cre* | F | Tamoxifen | 5 | 5 | 5 |  |

### Longitudinal Plasma Chemistry and NfL

Dietrich et al. (2017) reported that ubiquitous *Htt* loss leads to progressive neurodegeneration, including thalamic calcification (15). In hopes of determining when this process begins, we performed longitudinal plasma sampling from 5 animals per arm with matched WT controls to quantify the levels of NfL, a well-established marker of axonal stress, whose utility as a translational biomarker has been established in many neurodegenerative disease states (16). We collected plasma beginning at 3 months of age, continuing monthly until the study was complete at 14 months of age. In the *Htt^fl/fl^*;CAG-Cre line, we find that tamoxifen treatment leads to a rapid and persistent elevation of plasma NfL levels throughout the duration of the study (Fig 2A). At 3 months of age (1-month post-tamoxifen treatment), mean plasma NfL levels in tamoxifen-treated *Htt^fl/fl^*;CAG-Cre mice (2545 ± 678 pg/mL) are significantly increased compared to vehicle treated *Htt^fl/fl^*;CAG-Cre mice (840 ± 311 pg/ml; Tukey HSD p < 0.0001 ), both of which are higher than vehicle- or tamoxifen-treated *Htt^+/+^*mice, which are below 300 pg/mL (Fig. 2B; Tukey HSD p<0.0001). In the *Htt^fl/fl^*;UBC-Cre mice, we find a slightly different pattern. Initially, we *Htt^fl/fl^*;UBC-Cre are very similar to the *Htt^fl/fl^*;CAG-Cre mice – plasma NfL levels are significantly elevated at 3-months of age in tamoxifen treated mice (2091 ± 914 pg/mL), compared to vehicle (719 ± 165 pg/mL; Fig. 2B; Tukey HSD p < 0.0001). However, in the vehicle-treated *Htt^fl/fl^*;UBC-Cre mice, NfL levels increase steadily to 3467 ± 647 pg/mL, rising to levels similar to tamoxifen-treated mice (4430 ± 953 pg/mL) by 13-months of age (Fig. 2A). We hypothesize that this is driven by the much leakier nature of Cre in the CNS of the *Htt^fl/fl^*;UBC-Cre line, compared to the *Htt^fl/fl^*;CAG-Cre line (Fig. 1B), driving continued loss of HTT in the CNS during this period. From our final endpoint mice, we also conducted a cross-sectional screen of 11 plasma analytes and found that alanine transaminase (ALT), a biomarker of liver damage, was slightly elevated in tamoxifen-treated *Htt^fl/fl^*;UBC-Cre mice compared to vehicle (Fig. S6; Tukey HSD p=0.001).

**Figure 2.**
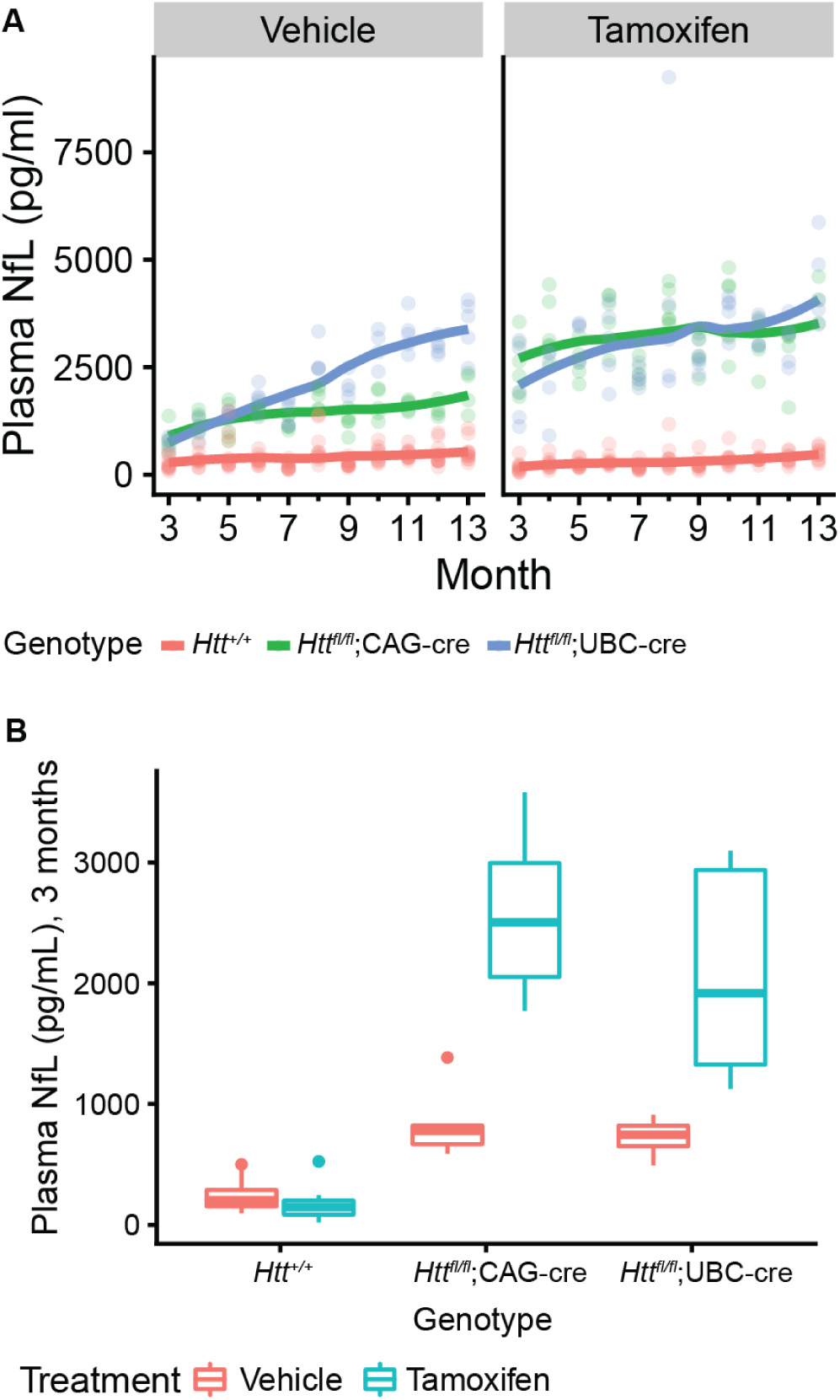
Plasma levels of neurofilament light chain (NfL) are increased after HTT loss. **A)** Plasma NfL levels are increased in all Htt^fl/fl^;CAG-Cre and Htt^fl/fl^;UBC-Cre mice, with the most significant increases being in the tamoxifen-treated groups. Plasma was collected once per month at 3-13 months of age and measured by chemiluminescent ELISA. **B)**. A cross-sectional view of NfL levels in each group at 3 months of age highlights the large increase in NfL after 1 month of HTT loss. n= 5 for cre groups, n=10 for non-cre groups (Htt ^+/+^ and Htt^fl/fl^ combined).

### Behavioral Analyses

Mice underwent a monthly modified SHIRPA exam to monitor neurological signs and several cross-sectional behavioral assays at 11 months of age (Table S1). We observed progressive tremor during the SHIRPA exam (Fig. 3A) in both cKO lines starting at 3 months and increasing in severity through 13 months. In conjunction, we observed limited and abnormal gait in some of the cKO mice, but this was not as consistent. All other SHRIPA measures appeared normal. For additional motor tasks, we excluded *Htt^fl/fl^*;UBC-Cre due to less controlled HTT knockdown in CNS tissue. As previously reported (15) we observed motor impairment, as revealed by an elevated balance beam traversal assay in which tamoxifen-treated *Htt^fl/fl^*;CAG-Cre mice take longer than vehicle-treated littermates (Fig 3B; *Htt^fl/fl^*;CAG-Cre tamoxifen vs. *Htt^fl/fl^*;CAG-Cre vehicle; Tukey HSD p < 0.0001). We also conducted an examination of the total movement using an open field assay, revealing a potential sustained impact of tamoxifen treatment, rather than HTT loss, as tamoxifen treatment of both *Htt^fl/fl^* and *Htt^fl/fl^*;CAG-Cre mice leads to modest hypoactivity (Fig. 3C; *Htt^fl/fl^* tamoxifen vs. *Htt^fl/fl^* vehicle Tukey HSD p<0.02; *Htt^fl/fl^*;CAG-Cre tamoxifen vs. *Htt^fl/fl^*;CAG-Cre vehicle Tukey HSD p<0.04). In summary, we found that HTT loss is associated with progressive neurological alterations, as revealed by increasingly severe tremors and motor impairment.

**Figure 3.**
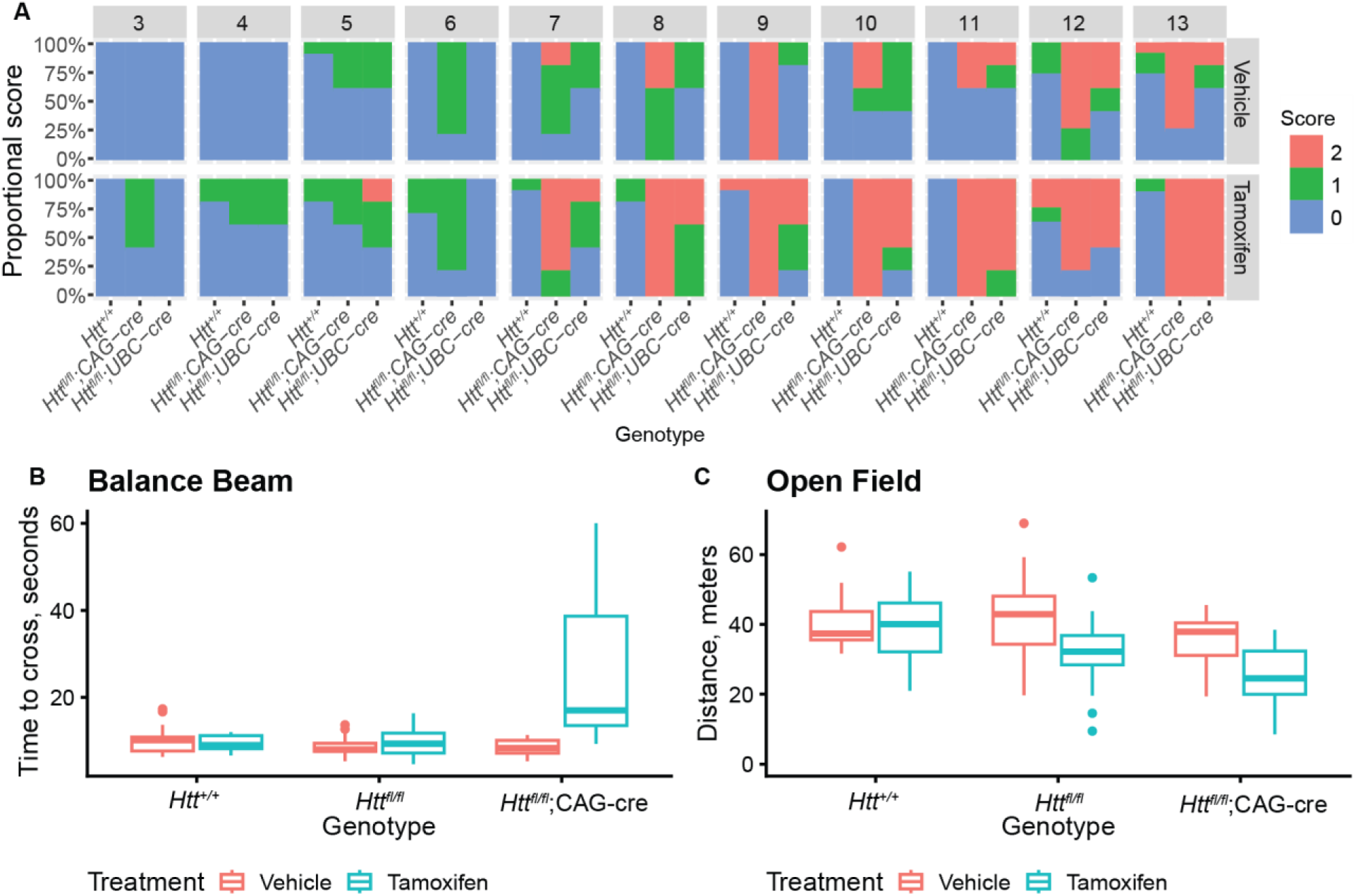
Progressive behavioral alterations develop after HTT loss. **A)** Tremor incidence and severity in tamoxifen treated Htt^fl/fl^;CAG-Cre and Htt^fl/fl^;UBC-Cre mice progressively worsens with age. n= 5 for cre groups, n=10 for non-cre groups (Htt ^+/+^ and Htt^fl/fl^ combined). **B)**. At 11 months of age, tamoxifen-treated Htt^fl/fl^;CAG-Cre mice take significantly longer (2.8x, p<0.001) to cross a balance beam than control mice. N = 15-20/group. **C)** At 11 months of age, tamoxifen-treated Htt^fl/fl^ and Htt^fl/fl^;CAG-Cre mice travel slightly less distance than control mice in an open field test, a result of possible tamoxifen toxicity. N = 15-20/group.

### Terminal HTT and NfL levels

Our interim analyses of HTT levels (Fig 1) and NfL (Fig 2) were based on relatively small numbers of animals. At 14-months of age we collected a more robust dataset from the complete endpoint cohort of mice, providing an opportunity to collect both cerebrospinal fluid (CSF) and plasma to directly compare NfL levels in both compartments. In general, our HTT assays reveled a pattern consistent with our interim HTT lowering data (compare Fig 1C and Fig 4A) – namely that the *Htt^fl/fl^*;CAG-Cre line was relatively well-regulated in the CNS, while the *Htt^fl/fl^*;UBC-Cre line was leaky in the CNS but more tightly regulated in the periphery (Fig. 1B). Similarly, both the CSF and plasma NfL quantifications revealed patterns consistent with those observed in our longitudinal study (compare Fig 2A and Fig 4B).

**Figure 4.**
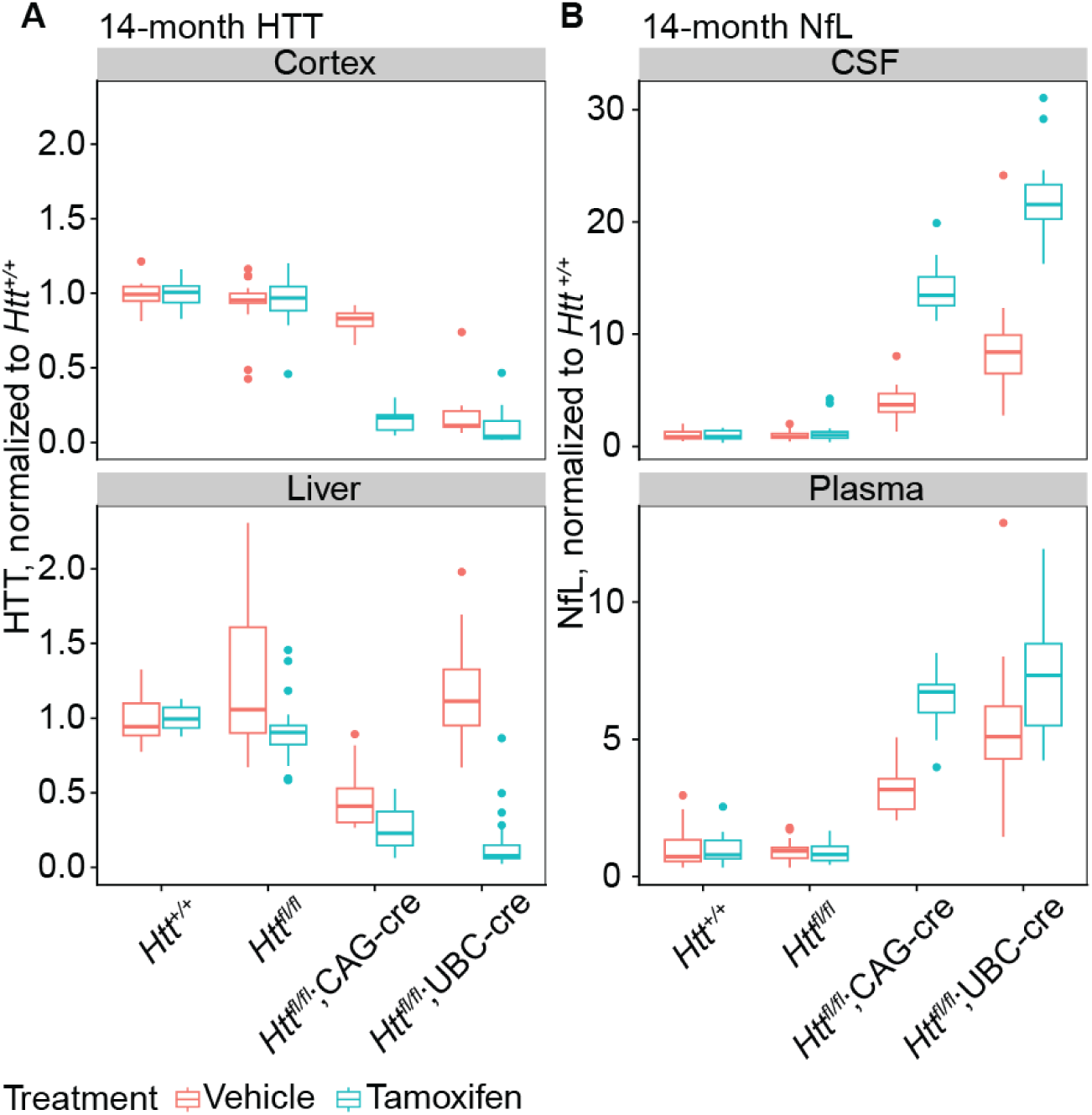
Terminal HTT levels are lowered and NfL levels are elevated in tamoxifen-treated cKO mice. **A)** In cortex (above), vehicle-treated Htt^fl/fl^;CAG-Cre mice have HTT levels similar to control groups. However, vehicle-treated Htt^fl/fl^;UBC-cre have reduced HTT levels similar to tamoxifen-treated mice, demonstrating leakiness of UBC-cre in the CNS. In liver, the inverse can be seen, demonstrating leakiness of CAG-cre in the periphery. In both tissues, tamoxifen-treated cKO mice show robust reduction of HTT. N = 12-20/group. **B)** NfL levels from CSF (above) and plasma (below) at 14 months of age show significant increases which correspond to the reduction of HTT seen in panel A. N = 11-20/group.

### Peripheral Pathology

Given that early loss of *Htt* before 3 months of age had been reported to cause fatal pancreatitis, we were particularly interested in the impact of HTT loss at 2 months of age on the pancreas. We examined the amount of HTT loss in the pancreas across our lines – as with other peripheral organs, we found that the *Htt^fl/fl^*;CAG-Cre mice were slightly leaky, while the *Htt^fl/fl^*;UBC-Cre line much less so (Fig. S1). After tamoxifen treatment, we observe significant lowering at 6 months (*Htt^fl/fl^*;CAG-Cre compared to *Htt^fl/fl^*: 93% reduction, Tukey HSD p = .002; *Htt^fl/fl^*;UBC-Cre compared to *Htt^fl/fl^*: 87% reduction, Tukey HSD p = 0.006), but the difference is not significant at the 3 or 9 month timepoints. This is likely because we had difficulty dissecting the pancreas without contamination by the surrounding adipose tissue, as well as the high level of proteases in pancreatic tissue causing high variability of HTT measurements compared to other tissues analyzed here (Fig. 1C). At 9 months of age, we sacrificed mice and collected pancreases for histological examination. Pancreatic anatomy in all genotypes and treatment groups was normal (Fig. 5A) and contained no diagnostic lesions. Pancreatic acinar cells are filled with zymogen granules and featured no signs of degeneration, necrosis, or inflammation, while islet cells were within normal limits. This provides more evidence that early HTT loss is not associated with symptomatic pancreatitis, pancreatic inflammation, or degeneration of the pancreas.

**Figure 5.**
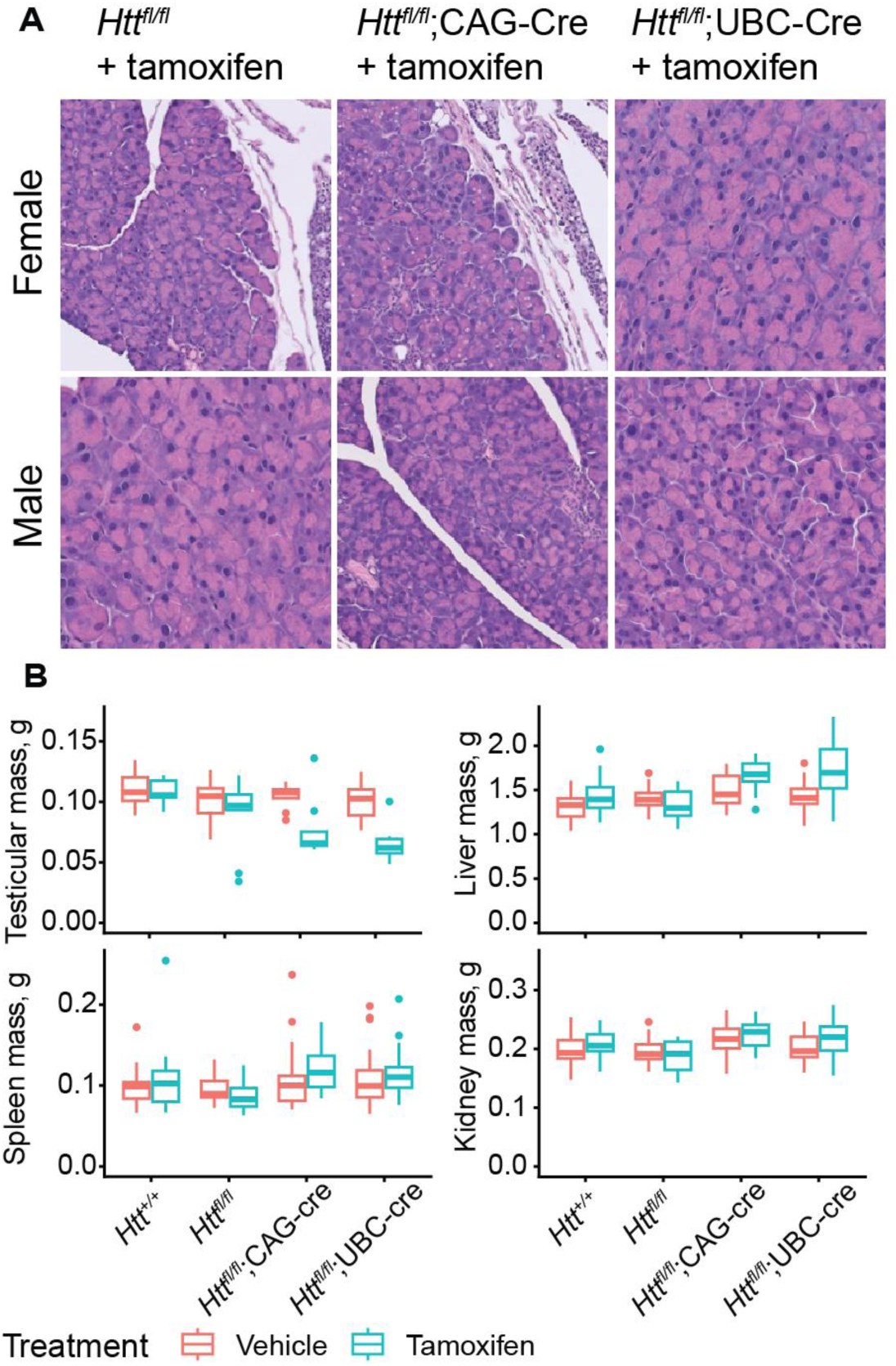
Pancreatic histology is normal after early and prolonged HTT loss. **A)** H&E staining in the pancreas shows no gross abnormalities in any genotype of tamoxifen treated mice. **B)** Testicular weights are lower in both tamoxifen-treated Cre lines compared to non-tamoxifen treated Cre lines. Other organ weights show no significant changes. N = 12-25/group.

We monitored body weight longitudinally and observed a complex relationship between sex, aging and HTT loss, but in general, alterations in bodyweight that may be associated with HTT loss are very subtle in our cohorts (Fig S2). Consistent with Dietrich et al. (2017), we observe loss of testicular mass in both Cre lines after tamoxifen treatment, though this didn’t reach statistical significance as part of our factorial ANOVA comparing all genotypes and treatments (ANOVA Treatment x Genotype interaction p = 0.65). If we restrict our focus on male mice of both Cre genotypes, the impact of tamoxifen treatment is robustly significant, supporting the validity of this reduction (Fig 5B, Treatment ANOVA main effect p < 0001). We recently reported that constitutive hepatic *Htt* knockout results in blistering of the Glisson’s capsule when the liver is perfused with PBS (21). We tested this here in a subset of mice at 14-months and observed the same blistering in 100% of cKO mice tested (5/5 tamoxifen treated *Htt^fl/fl^*;UBC-Cre, 0/5 of *Htt^fl/fl^*, Fisher’s Exact p = 0.0079), suggesting this loss of adhesion is replicated after adult-onset HTT loss.

### Central Pathology

Consistent with Dietrich et al (2017), we observe very large subcortical lesions surrounding accumulations of calcified deposits in Cre mice treated with tamoxifen. In a sagittal section stained with a calcium-sensitive dye, Alizarin Red, a very clearly demarcated thalamic lesion becomes clear (Fig. 6A). Manual alignment of each mouse’s lesions to the Allen Institute reference atlas(22) reveals a relatively circumscribed localization to the posterior complex, ventral posteromedial nucleus, and ventral anterior-lateral complex of the thalamus, which comprise the sensory nuclei of the thalamus, as well as less frequent lesions found in the mediodorsal, paracentral, and ventral posterolateral nuclei of the thalamus (Fig. 6B). Calcifications are also clearly visible in sections stained with Cresyl violet (Fig. S4). To measure the atomic composition of the lesions more objectively, we turned to scanning electron microscopy-energy-dispersive X-ray spectroscopy (SEM-EDS), which can determine the elemental composition of a sample by characterizing the X-rays emitted as the electron beam sweeps across the sample (23). Paired comparison of spots within and outside the lesion reveals a clear shift – the signal from non-deposit tissue is dominated by carbon, with smaller peaks of calcium, oxygen, and minor peaks for other elements (including silicon, from the underlying glass slide; Fig. 6D). This profile is markedly different within the lesions, which are depleted of carbon, with a strong increase in the calcium and phosphate peaks. We do not observe any significant peaks at 6.4 keV, the characteristic energy level of Iron, suggesting that the deposits found in our HTT depleted mice are not notably enriched in iron above the lower limit of detection with this technique. Western blots for transferrin receptor (TFRC) were conducted to assay changes in iron regulatory proteins found in Dietrich et al. (2017). TFRC is a key iron import protein and is upregulated when intracellular iron levels are low (24). We found that TFRC was significantly increased in tamoxifen-treated *Htt^fl/fl^*;CAG-Cre mice compared to tamoxifen-treated *Htt^fl/fl^* mice (Fig. 6C, Welch’s T test p < 0.002).

**Figure 6.**
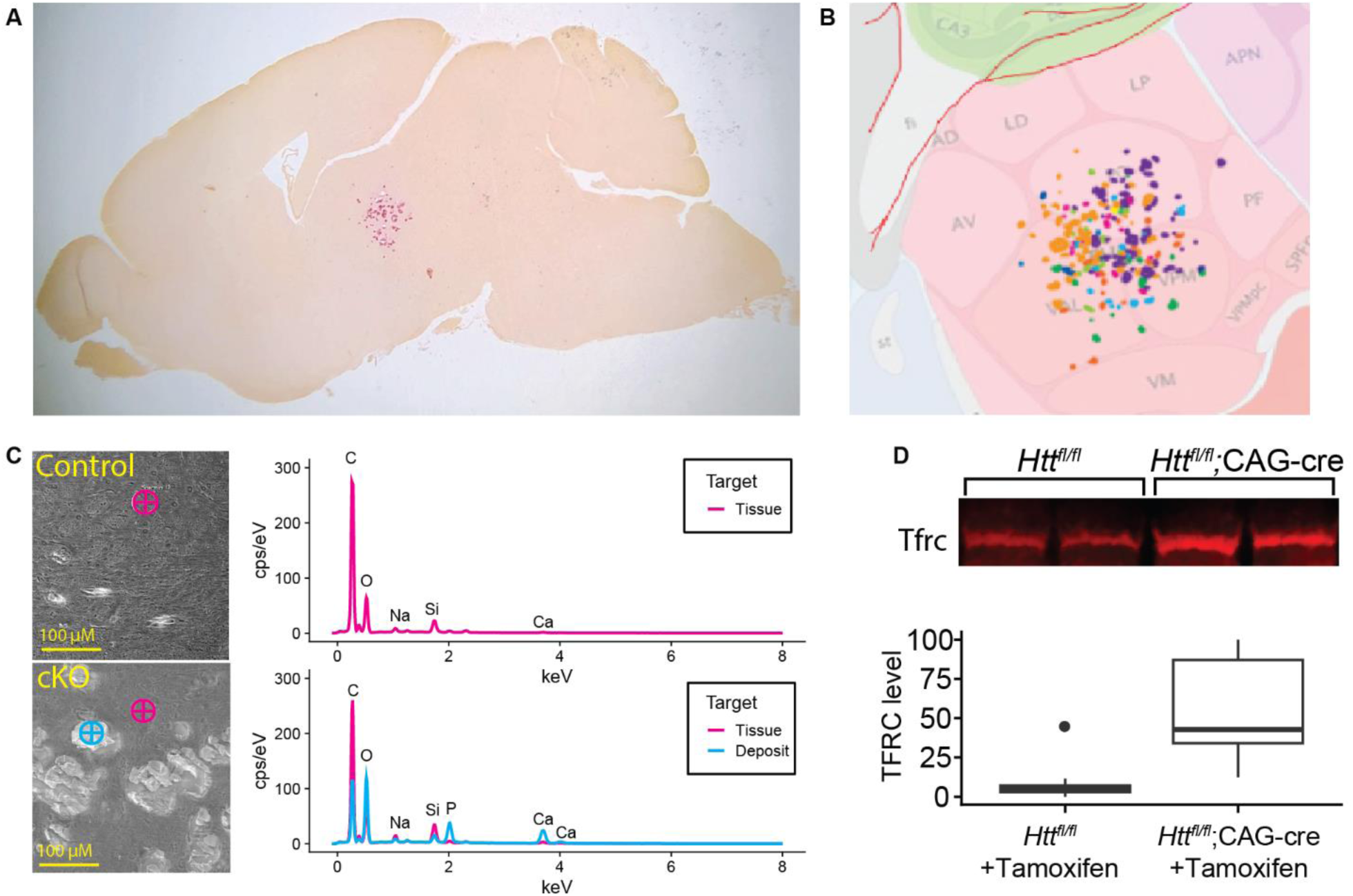
Histological analysis reveals overt thalamic calcification and inflammation after HTT loss. **A)**. Large subcortical deposits are strongly positive for calcium marker Alizarin red. **B)** Overlay of Allen Brain Atlas on lesions demonstrates the average location of the lesions, where lesions from every mouse are represented by a different color (see Fig. S3 for more detail). **C)** (Left) SEM-EDS images reveal that deposits are absent from control tissue but present in cKO tissues. (Right) EDS spectra reveals that deposits show enrichment for oxygen (O), and phosphate (P), and calcium (Ca) suggesting these are likely Ca2PO4 deposits. **D)** Western blot of cortical lysates reveals increased levels of Tfrc in cKO mice; Tfrc signal was normalized to total protein per well.

We collected sagittal sections of all mice in our study for the examination of histological outcomes. Sections were stained with antibodies for neurons (NeuN, a pan-neuronal marker), astrocytes (GFAP) and microglia (ionized calcium-binding adaptor molecule 1; IBA1) (Fig. 7A). Thalamic GFAP intensity is significantly higher in tamoxifen-treated *Htt^fl/fl^*;CAG-Cre mice than in vehicle-treated or wildtype mice (Fig. 7B, Tukey HSD p = 0.0001), supporting findings by Dietrich et al (2017). Additionally, IBA1 positive cells appear more numerous and surround the calcified lesions in tamoxifen-treated *Htt^fl/fl^*;CAG-Cre sections, though quantified IBA1 positive area was not significantly higher overall (ANOVA Treatment x Genotype interaction p = 0.057).

**Figure 7.**
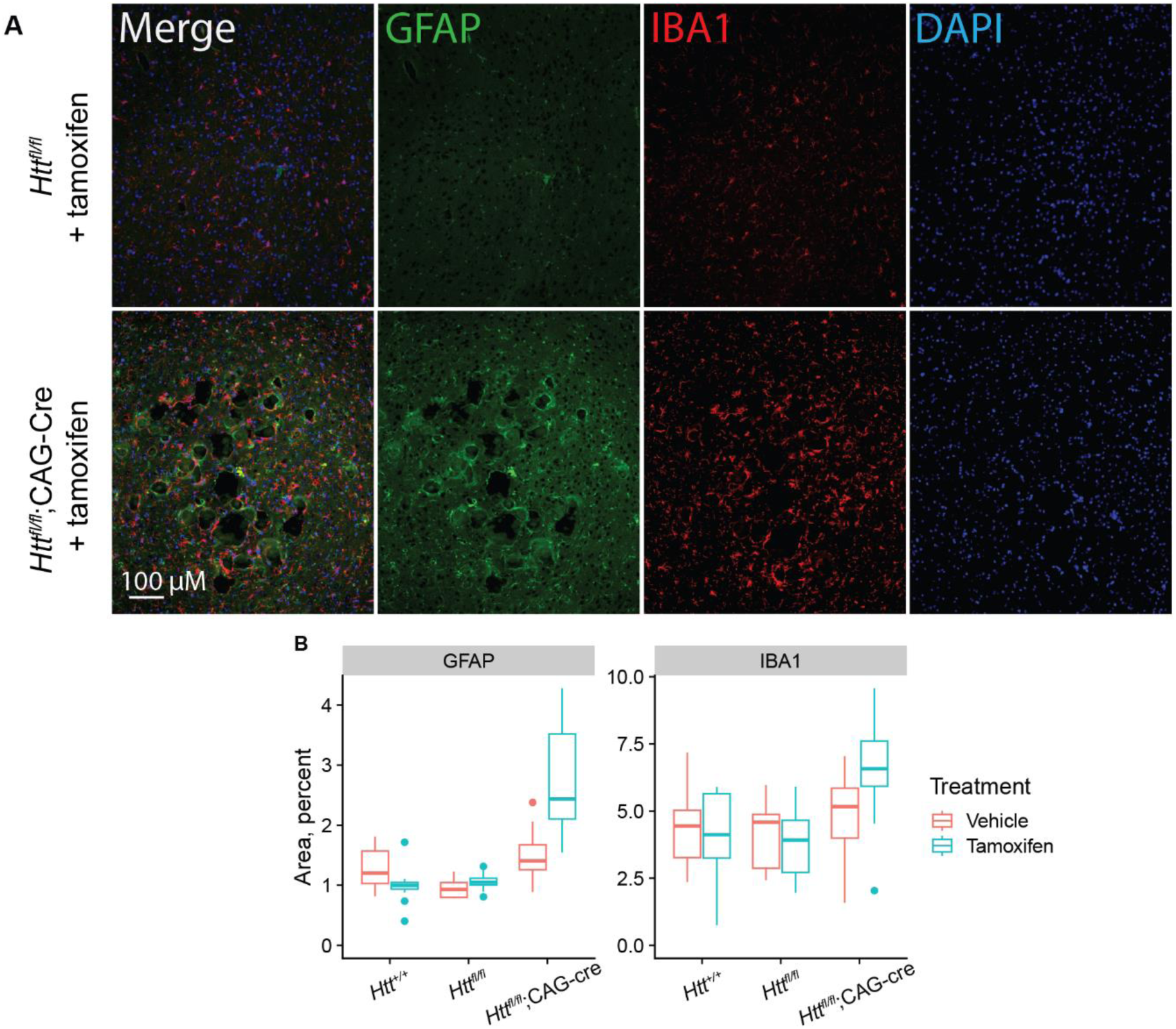
Immunoreactivity for glial markers IBA1 and GFAP is increased after HTT loss. **A)** Immunohistochemical staining for microglia (IBA1, red) and astrocytes (GFAP, green), nuclei (DAPI, blue) reveal inflammation in the thalamus surrounding the calcified deposits. **B)** Quantification of IBA1 and GFAP area reveal significant increase of GFAP area in cKO mice, while IBA1 which qualitatively appear increased surrounding the thalamic lesions, is not significantly increased in cKO thalamus overall.

## Discussion

This study was conducted to replicate an apparently discrepant set of results between two published studies of the impact of global adult-onset *Htt* knockout. We find that our results largely support those of Dietrich et al (2017) (15) and further reveal significant discrepancies with Wang et al (2016) (14). While we don’t fully understand the origin of these differences, some possible explanations are discussed below. In short, we confirm that the chronic, near-complete cellular loss of HTT at 2 months of age is initially benign, but associated with a progressive neurodegenerative phenotype accompanied by increased levels of NfL in the plasma. We also describe the results of SEM-EDS work confirming that the primarily thalamic lesions are calcium phosphate rich. This constellation of signs raises cautions about a lower limit of HTT silencing that is tolerated *in vivo*, particularly in the brain. Future studies with more graded *Htt* lowering are justified to determine whether less extreme lowering carries any of the same risks.

The impact and tolerability of HTT loss in adulthood has been a controversial and complex issue for some time (see recent reviews on the issue (25,26)). It is now clear that complete loss of HTT, or expression of hypomorphic alleles, is not tolerated during development in mice (2–4,27), or humans (5,6). However, graded reductions of HTT mediated by small interfering RNA (siRNA) approaches in both mice (28) and non-human primates for as long as six months have been reported to be benign (29–33). Similarly, HD patients treated with Tominersen and other CNS-directed HTT-lowering drugs have not been reported to develop subcortical calcification. However, many patients treated with 120mg doses of Tominersen, which resulted in robust lowering of HTT in the CSF, did show a puzzling, transient increase in NfL levels—the opposite of what was predicted at study onset (12). It may be that the increased CSF NfL in HD patients treated with Tominersen shares a mechanistic origin with the large and sustained increases in plasma NfL we observe after near-complete HTT loss in mice, though unlike the human trial we did not observe a return to baseline in NfL throughout the 12 months of our study.

A major motivation for this study was to clarify whether complete HTT loss is safe, in hopes of informing the rapidly moving field of HTT lowering in HD clinical trials (8). We find that near-complete loss of HTT is poorly tolerated in the brain, suggesting that a straightforward genome editing approach to complete elimination of both HTT alleles is likely not a desirable approach to HD therapeutics (34) and that the allele-selective genome editing approaches that have been proposed would be better tolerated (35–37). However, an important limitation of this study is that we used an experimental tool – tamoxifen-inducible Cre – that drives HTT cellular expression from normal levels to near-zero very quickly, which is distinct from pharmacological approaches such as ASOs which result in graded lowering of target genes across cell types (38). The ubiquitous and strong promoters used in this study also drive expression across every cell type of the body (and brain), and so deconvolving the impact of HTT loss on different cell types is not feasible with our approach. Developmental deletion of HTT in specific cell lineages has been shown to be deleterious via distinct mechanisms – e.g. HTT loss in postnatal neurons and testes results in progressive neurodegeneration and impaired spermatogenesis (39); and loss of *Htt* in *Wnt1*-lineage cells is associated with congenital hydrocephalus (40).

Given that this study was conducted primarily as a replication study, we need to consider points of coherence and points of discrepancy between the now 3 studies conducted to address the safety of global inducible *Htt* loss. The mice Dietrich et al (2017) relied on were *Htt^fl/-^*;CAG-Cre (referred in that study as “cKO”), meaning that they developed with 50% of normal total HTT levels before being treated with tamoxifen to drive HTT from 50% towards 0%. In general, we observe very few differences from the findings of this study, apart from body weight, which they observe is consistently reduced in tamoxifen treated “cKO” mice, while we observe a more complex relationship between age, genotypes and sex (Fig. S2). This may be due to the confound of different total HTT levels during development in their study (50%) and ours (100%). Interestingly, Dietrich et al (2017) report some unexpected phenotypes in cKO mice not treated with tamoxifen, namely: earlier death, lower body weight (consistent with a previous finding (13)) and impaired motor coordination compared to wildtype mice. This suggests that 50% HTT expression is associated with significant important age-related phenotypes, which were likely exacerbated by additional loss of HTT expression that we observe occurs with CAG-cre and the Htt flox allele.

Our findings are more discrepant with those of Wang et al (2016), who observed acute pancreatitis after treating *Htt^fl/fl^*;CAG-Cre at 2 months of age with tamoxifen, but no deleterious motor or pathological phenotypes with mice aged to 18 months of age after knocking out HTT at 4 or 8 months of age. We speculate that this apparent discrepancy may arise from selective toxicity of tamoxifen, a compound with a very narrow therapeutic index in both humans and mouse models (41), particularly in 2-month-old animals. The levels of tamoxifen injected by the Wang et al (2016) were 100 mg/kg for 5 consecutive days, while we and Dietrich et. al (2017) used 75-77 mg/kg. While the higher tamoxifen levels used by Wang et al (2016) may explain the early toxicity observed, it remains unclear why *Htt^fl/fl^*;CAG-Cre mice would be selectively vulnerable to this compared to non-floxed control mice, who survived. The apparently benign mouse phenotype observed by Wang et al (2016) after knocking out HTT at 4 and 8 months of age may reflect the reduced amount of time those mice experienced acute HTT loss before sacrifice, though we observe clear behavioral outcomes such as tremors as early as 5 months post HTT loss (Fig. 3A). It may be the case that silencing HTT at 2 months of age is particularly deleterious because of the developmental stage at which silencing occurs.

We are intrigued that complete HTT loss reproducibly causes subcortical calcification, a very distinctive neuropathological lesion. Many human conditions, and some of their animal models, have similar lesions. One such family of diseases—type I interferonopathies—is caused by hyperactivation of type I interferon responses secondary to deficits in nucleic acid homeostasis (42,43). Humans with these conditions often present with subcortical calcification, as do many of the mouse models generated to study them. Similarly, patients with Cockayne syndrome – which arises from deficient DNA repair – often present with subcortical calcification (44). In Down syndrome, calcification of the basal ganglia has been widely observed and has been attributed to enhanced brain aging (45,46). Another potentially significant point of comparison is with familial idiopathic basal ganglia calcification (FIBGC; Fahr’s disease) which is a common cause of intracranial calcification in humans (47) An emerging picture of the pathology in FIBGC is that it arises from deficiencies in ion homeostasis across the blood-brain barrier (BBB), either via mutations in ion transporters themselves (SLC20A2 (48) or XPR1 (49)), or in genes that play a role in pericyte growth factor responses (PDGFB (50) or PDGFRβ (51)), and thereby BBB permeability. In recently proposed models, the high local calcium and phosphate concentrations drive deposition of CaPO4 deposits near tissues which produce CSF, which drives the anatomic localization of these highly localized deposits (47). Future detailed work comparing the phenotypes of mice with acute loss of *Htt* and these other syndromes associated with subcortical calcification is warranted by the reproducibility and specificity of this phenotype.

## Methods

### Mice

We generated two lines of conditional *Htt* knockout mice (cKO) by crossing CAG-Cre (JAX stock 004682) or UBC-Cre (JAX stock 007001) to Htt^fl^ mouse line (4). Mice were maintained at homozygosity for the Htt fl allele and heterozygosity for the cre allele. All mice were bred at the McLaughlin Research Institute (MRI) and housed in cages of 3-5 mice with access to food and water *ad libitum* unless otherwise mentioned. Vivarium lights were on a 12-hour light/dark cycle. The MRI institutional animal care and use committee approved all procedures under protocol 2020-JC-29. Mice were genotyped with the following primers: for Htt flox PGK Reverse 5’ cta aag cgc atg ctc cag act g 3’, Flank Forward 5’aga tct ctg agt tat agg tca gc 3’, and Flank Reverse 5’ cat ttg att ctt aca ggt agc ctg 3’ (band sizes of 320 bp for the Floxed PGK Reverse and Flank Forward and 180 bp for the WT with Flank Forward and Flank Reverse), for UBC-cre 25285 Forward 5’gac gtc acc cgt tct gtt g 3’, oIMR9074 Reverse 5’agg caa att ttg gtg tac gg 3’ (band size 475 bp), and for CAG-cre oIMR5984 Forward 5’ gct aac cat gtt cat gcc ttc 3’, oIMR9074 5’ agg caa att ttg gtg tac gg3’ (band size of 180 bp). Band sizes were determined on a 1.5 % agarose gel with 0.5% TBE buffer. Tamoxifen (Sigma T5648) was administered via I.P. injection at 75mg/kg once daily for 5 consecutive days. Tamoxifen was reconstituted in pharmaceutical grade corn oil (Sigma C8267) prior to administration.

### Recombination Analysis

DNA recombination was tested using semi-quantitative PCR (Fig. S5). Genomic DNA was extracted from cortex and liver using the IBI gMax Mini Kit (IB47281), using the manufacturer’s protocol for solid tissue. PCR was performed using the pgk, Hdhpr13, and Hdhrec9 primers flanking the deletion site as described by Dietrich et al. (15), resulting in two bands, a 220 bp one showing the amount of recombined allele and a 210 bp one showing the amount of unrecombined allele. PCR products were run on 2.2% agarose gels (Lonza 57031) and imaged using Philips VLounge software. Band intensity was quantified in ImageStudio Lite software. Recombination efficiency was calculated by dividing the intensity of the recombined band by the total signal of the recombined plus unrecombined bands, so that a higher value is equivalent to more recombination.

### Behavioral Analysis

Mice were assessed monthly beginning at 3 months until 13 months of as part of a modified SmithKline Beecham, Harwell, Imperial College, Royal London Hospital Phenotype Assessment (SHIRPA)(18,52). Briefly, mice were assessed for 13 measures (Table S1) where normal behavior received a score of “0” and severity of abnormalities were indicated with higher scores ranging from 1-3, depending on the measure as outlined on Table 4. To assess thigmotaxis behavior, mice were recorded for 10-minutes in an open field box (40 x 40 cm) and activity was quantified with ANY-Maze software (Stoelting) for measures including distance traveled and time spent within 6.5 cm of walls. The following two days, mice were trained to cross a 1-meter long elevated balance beam and traversal time was measured on the third day as described (53). For training, mice were placed on one end of the beam, which had a 60-watt lamp, and encouraged to cross the beam towards the opposite side, which had a dark box with home-cage nesting material. Mice were trained with three trials on a 12-mm wide beam with 15 second intervals between trials. Mice were then given a 10-minute break and this was repeated on a 6-mm wide beam. After two days of training, mice were similarly placed at the start of the 6-mm beam and the time to traverse was measured and averaged for three trials. Any time over 60 seconds was scored as 60 seconds, any falls were scored as 60 seconds.

### Protein Quantification (HTT, NfL, Tfrc)

Total HTT levels were quantified using an electro-chemiluminescent ELISA measured with a MESO QuickPlex SQ120MM (Meso Scale Discovery; MSD) according to previously described methods (54). Tissues were homogenized in non-denaturing lysis buffer in tubes containing 1.4mm zirconium oxide beads at 6 m/s in three 30 second intervals with 5 minutes on ice between rounds. Lysates were centrifuged for 20 minutes at 20,000 x g at 4C. Supernatant was transferred to a new tube and centrifuged again for 20 minutes at 20,000 x g at 4C, then transferred to a second new tube. 96-well MSD plates (MSD, L15XA-6) were coated with capture antibody (CHDI-90002133 , 8ug/mL) in carbonate-bicarbonate coating buffer for 1 hour with shaking at 750 RPM. Plates were then washed 3 times with wash solution (0.2% Tween-20 in PBS) and blocked with blocking buffer (2% BSA, 0.2% Tween-20 in PBS) for 1 hour with shaking, then washed 3 more times. Samples were diluted (brain: 2ug/uL, liver: 4ug/uL) in 20% MSD lysis buffer, 80% blocking buffer and incubated for 1 hour with shaking. Plates were washed 3 more times, followed by incubation with sulfo-tag conjugated secondary antibody (D7F7-Sulfo-Tag, 1:1800) for 1 hour with shaking. Plates were washed a final 3 times, and Read Buffer B (MSD, R60AM-4) was added to the plate before reading on a QuickPlex SQ 120MM.

Neurofilament light levels were quantified using the Neurofilament L Assay (MSD, K1517XR-2) according to manufacturer directions. Briefly, plates were coated with biotinylated capture antibody at 1:16.5 in Diluent 100 and incubated for 1 hour with shaking at 700 RPM. They were then washed 3 times with wash buffer, followed by addition of the samples diluted at 1:10 in Diluent 12, and incubated for 1 hour at 700 RPM. After 3 more wash steps, secondary antibody was added at 1:100 in Diluent 11 and incubated for 1 hour at 700 RPM, followed by a final 3 washes. Read Buffer B was added, and the plate was read on a QuickPlex SQ 120MM.

TFRC was quantified by Western blot. Samples were run on a 10% Bis-tris gel (Fisher, NO0301) for 40 minutes at 200V. The iBlot 2 system with a PVDF membrane was used for protein transfer. Membranes were blocked with Intercept blocking buffer (Licor 927-60001) for one hour, and incubated with TFRC primary antibody (Fisher, 13-6800) for one hour, followed by washing and then secondary antibody (Licor, 926-68070) for one hour. Total protein per lane was quantified using Revert total protein stain (Li-Cor) and each lane was normalized to the total protein per lane. Blots were imaged on a Li-Cor Odyssey CLx and quantified using ImageStudio (Li-Cor)

### Plasma Chemistry

Clinical chemistry plasma analysis for 11 analytes (Fig S3) was completed using an Atellica Clinical Chemistry Analyzer (Siemens) at Phoenix Central Laboratories (Mukilteo, WA). Plasma was collected via cardiac puncture following a lethal injection of sodium pentobarbital (Fatal Plus, Henry Schein) at 14 months (SD = 5.7 days). Plasma was flash frozen following collection in heparinized microtainers (BD, cat. no. 365965) and purification by centrifugation at 1,300 x g for 10 minutes, followed by 2,500 x g for 15 min.

### Histology

Hemibrains were drop-fixed in 10% neutral buffered formalin (BBC biochemical) for 48 hr. After fixation, they were stored in PBS + 0.02% sodium azide until they were paraffin embedded, cut into 5-μm sections, and mounted on glass slides. For fluorescent immunohistochemistry, sections were deparaffinized and rehydrated to distilled water as described previously [67]. Sections underwent heat-mediated antigen retrieval in a water bath for 20 min in pH = 9 Tris-EDTA buffer. After washing with water, slides were blocked for 1 hour with 20% goat serum in PBS. The primary antibodies were incubated at the following concentrations in PBS with 20% goat serum, 1% BSA, and 0.1% Tw20: GFAP (EMD MAB3402, 1:500), Iba1 (Wako 019-19741, 1:500), NeuN (EMD ABN78, 1:750). Samples were washed with PBS three times and incubated with the appropriate secondary antibody (Alexa Fluor) diluted 1:1000 for 1 hour at room temperature in antibody diluent. Samples were washed with PBS three times and coverslips were applied with Vectashield DAPI hardset mounting medium (Vector labs, H-1500-10). Slides were imaged on a Leica DMI600 widefield microscope. Quantification was completed in FIJI/ImageJ by thresholding to select image areas positive for immunostaining.

For calcium staining, sections were deparaffinized and rehydrated to distilled water as described previously. Sections were stained with a 2% Alizarin red solution (C.I. 58005) with pH adjusted to 4.2 using ammonium hydroxide, then rinsed in water. They were dehydrated with 20 dips in 100% acetone, then 20 dips of 1:1 acetone/xylene and cleared in 100% xylenes. Coverslips were applied with Permount (Fisher Chemical SP15). Slides were imaged on an Olympus BX51. Quantification was completed using a visual rating of “strong”, “weak”, or “no” staining while blinded to genotype and treatment. For pancreatic pathology, H&E-stained pancreatic sections were scored by a veterinary pathologist (Zoetis Reference Laboratories, Mukilteo, WA) blinded to genotype.

### SEM-EDS

Brains from 12-month-old mouse were drop-fixed in 10% neutral buffered formalin (BBC biochemical) for 48 hours followed by cryoprotection by sinking in 30% sucrose for 48 hours and subsequently flash frozen in isopentane on dry ice. Brains were embedded in cutting medium (OCT) and cut in 20um sections on a cryostat. Brains were mounted onto slides and allowed to dry overnight and coated in Au:Pd prior to imaging. Slides were imaged on a FEI Sirion XL30 Scanning Electron Microscope (SEM) equipped with an Oxford Instruments Energy Dispersive X-ray Spectrometer (EDS) under vacuum conditions. EDS spectra were collected from lesion areas and adjacent non-lesioned tissue or equivalent region of non-lesioned brains.

### Statistical analysis

Primary data for our analyses were processed using R [69]. For factorial tests, we used ANVOA with Tukey’s tests for post hoc analyses where appropriate. Data presented in Figs 1-7, S1, S2, and S5 used boxplots - horizontal lines indicate 25th, 50th and 75th percentile, while the vertical whiskers indicate the range of data. Data falling outside 1.5 times the interquartile range are graphed as isolated points but were not excluded from statistical analysis. Graphics were produced using ggplot2 [70], Illustrator (Adobe), and BioRender.com.

## Data Availability

All raw data underlying these analyses, including tabular molecular and behavioral data, R scripts and microscopy images are available as a stable digital repository at Dryad at with the persistent DOI: 10.5061/dryad.9zw3r22kt.

## Acknowledgements

Funding for this project was provided via an unrestricted research grant from Ionis Pharmaceuticals and research agreement between CHDI foundation and UW and WWU (Record number # A-18222). The authors thank Holly Kordasiewicz for thoughtful discussions around the design of this study. The authors thank Scott Braswell for assistance with SEM-EDS at the University of Washington Molecular Analysis Facility, a National Nanotechnology Coordinated Infrastructure (NNCI) site which has partial support from the National Science Foundation via awards NNCI-1542101 and NNCI-2025489. The authors thank the staff at McLaughlin Research Institute that assisted with behavioral assays and tissue collection including Serena McElroy, Megan Ratz-Mitchem, June Pounder, Kaela Davey.

## Supplemental Figures and Tables

**Supplemental Figure S1:**
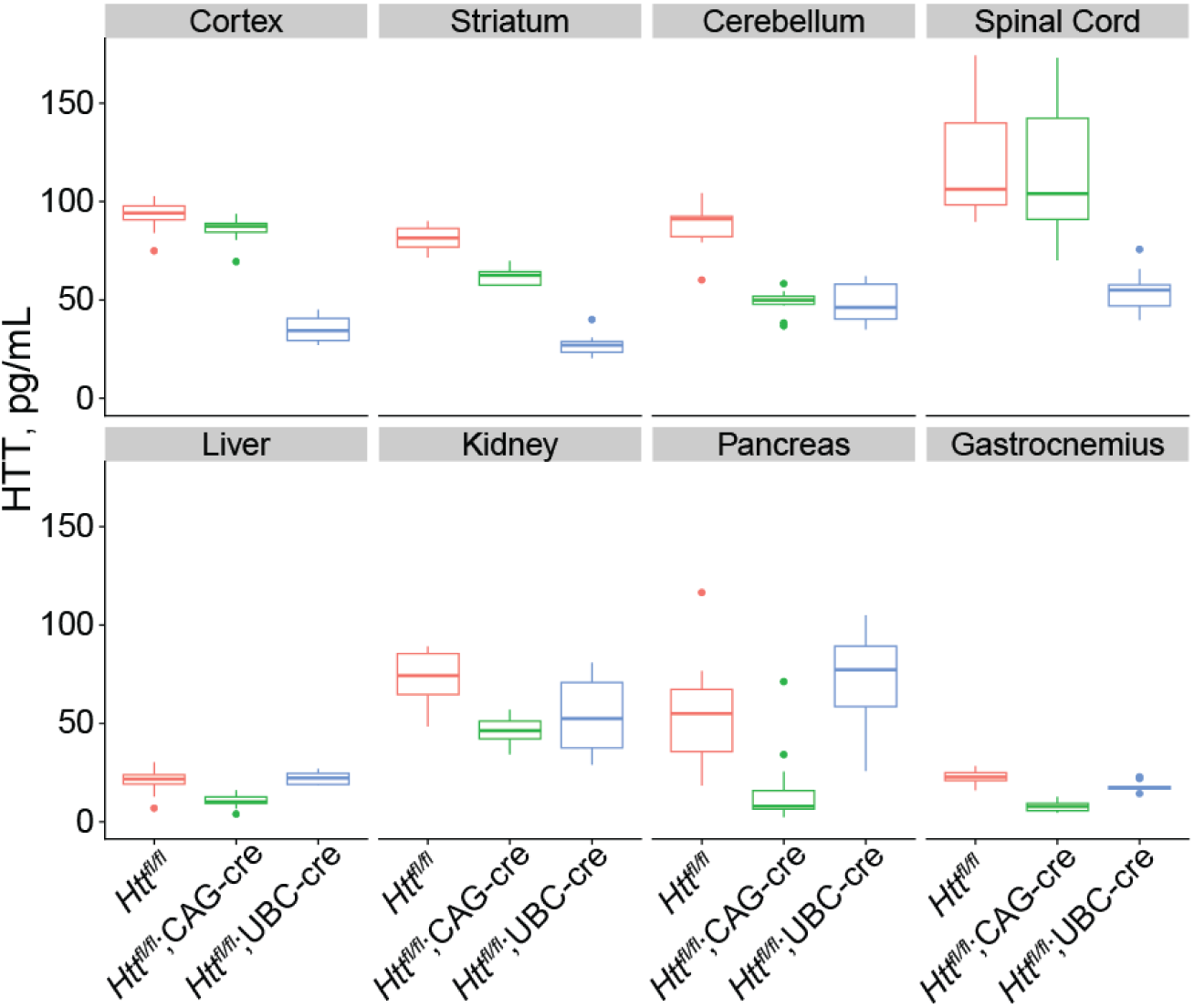
Quantification of HTT levels in the absence of tamoxifen in a wide range of tissues at 5 months of age. In the absence of tamoxifen, HTT levels in cKO and floxed control mice were measured in a variety of tissues. Peripheral tissues followed the same pattern we observed in the liver: reduced HTT in CAG-Cre mice, indicating leakiness, but relatively preserved HTT levels in UBC-Cre mice. Most brain tissues followed the opposite pattern, with the exception of cerebellum, which appeared leaky in both cKO lines. N = 10-12/group.

**Supplemental Figure S2:**
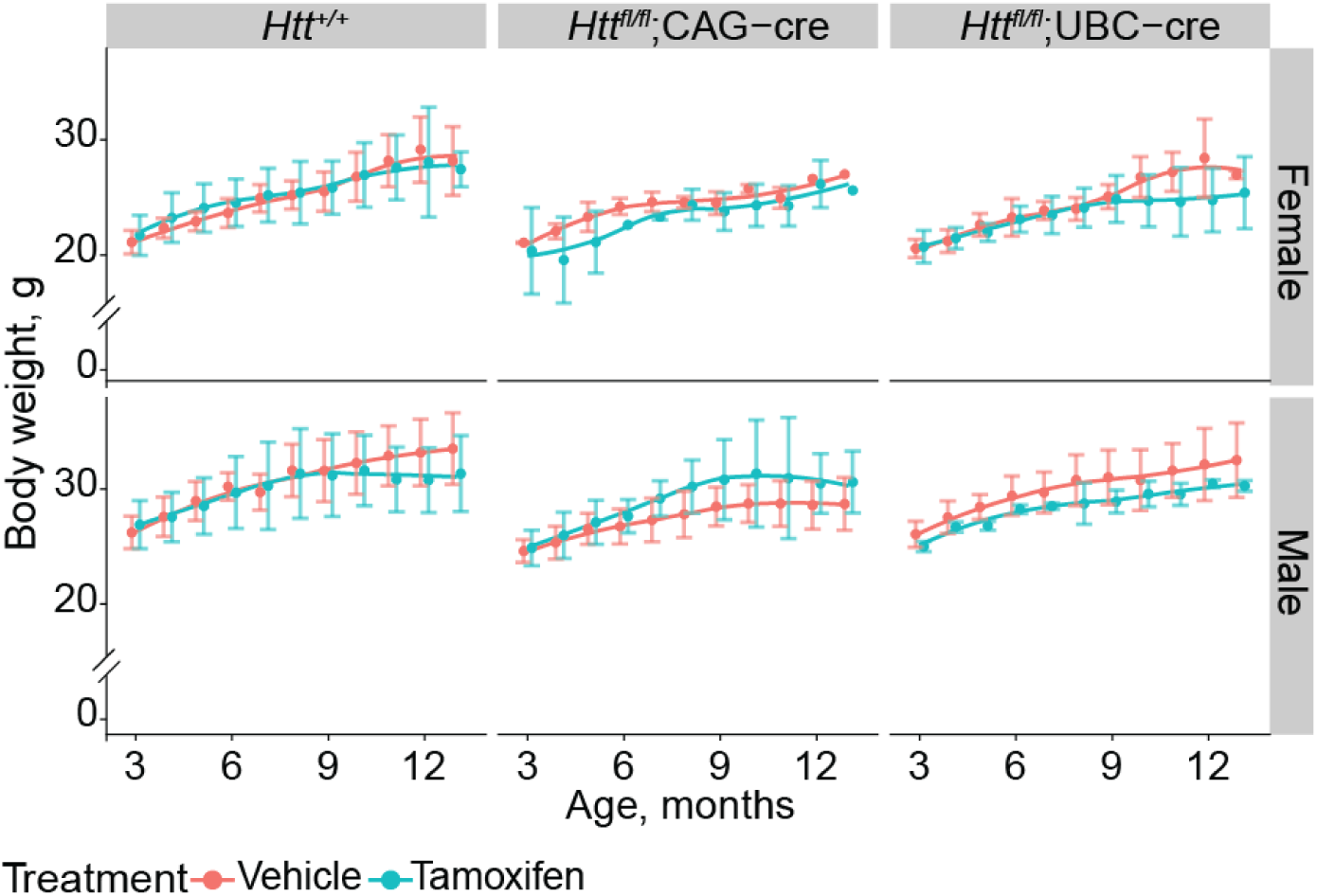
Longitudinal Body Weight after HTT loss. We monitored body weight over the lifespan of a subset of mice over their lifespan and observe very subtle differences across age and treatment. N=5-10 per group. *Htt^+/+^* and *Htt^fl/fl^* mice are pooled for visualization.

**Supplemental figure S3:**
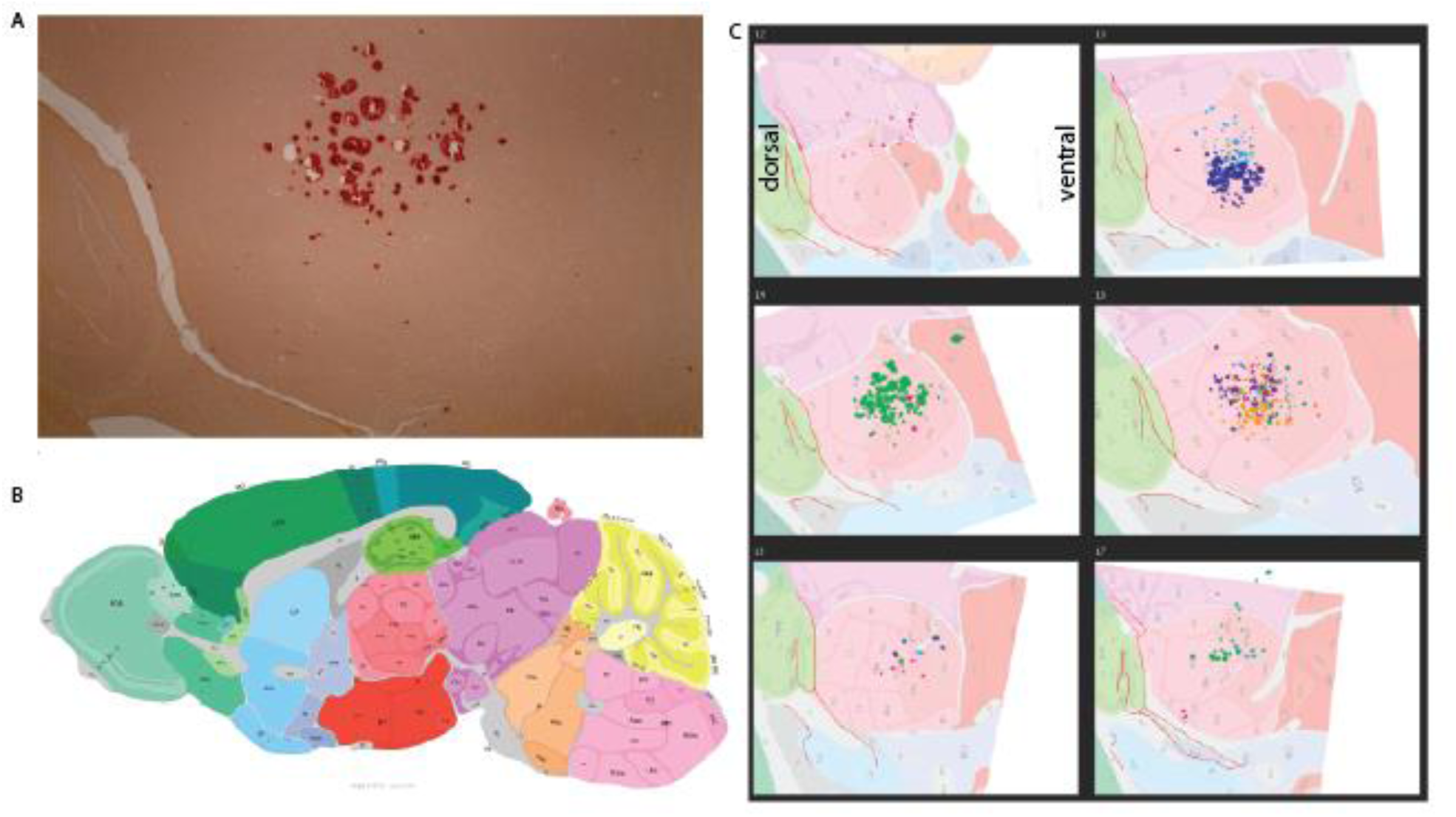
The precise location of the calcification was determined using the Allen Mouse Brain Atlas. Brain sections stained with Alizarin red (A) were categorized into a frame of the Allen Mouse Brain Atlas (B) depending on their sagittal position. All sections fell between frames 12 (most lateral) and 17 (most medial). For each frame number, images of the calcified areas were aligned with the corresponding atlas images based on the positions of the dentate gyrus and lateral ventricle. Areas of calcification were then traced onto the brain atlas image (C) and the anatomical positions were recorded. Color represents separate mice.

**Supplemental figure S4:**
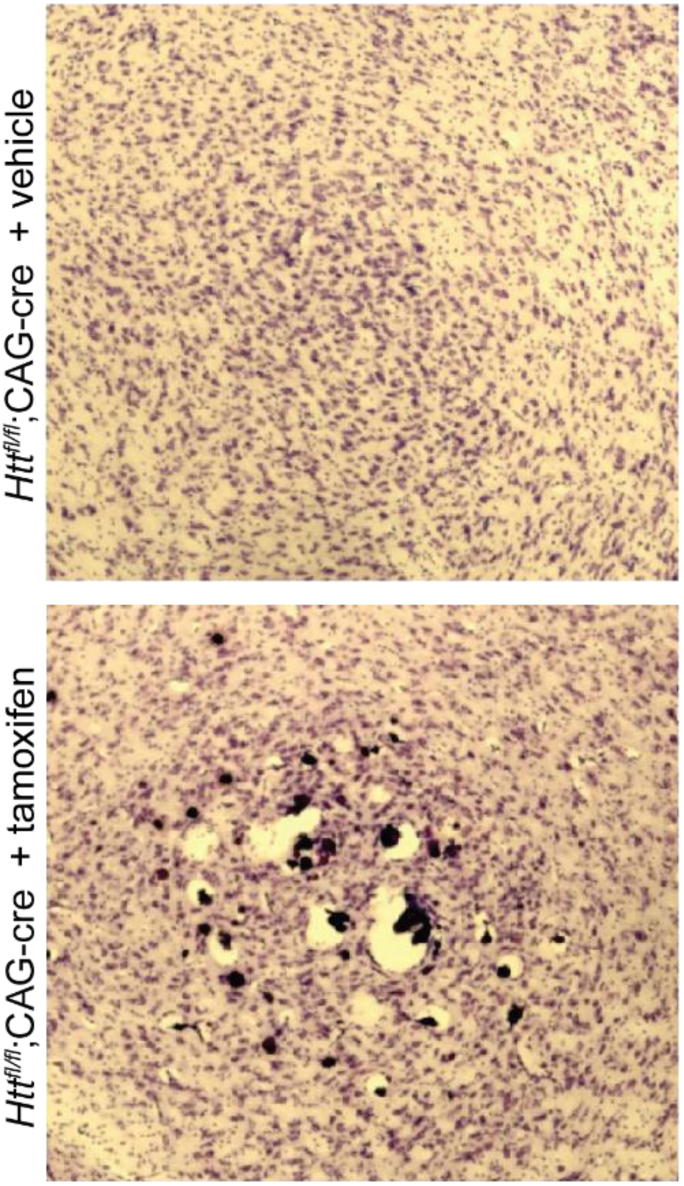
Thalamic calcification is visible in Cresyl violet staining. Sagittal brain sections were collected from cKO mice and stained with Cresyl violet to highlight lesions.

**Supplemental figure S5:**
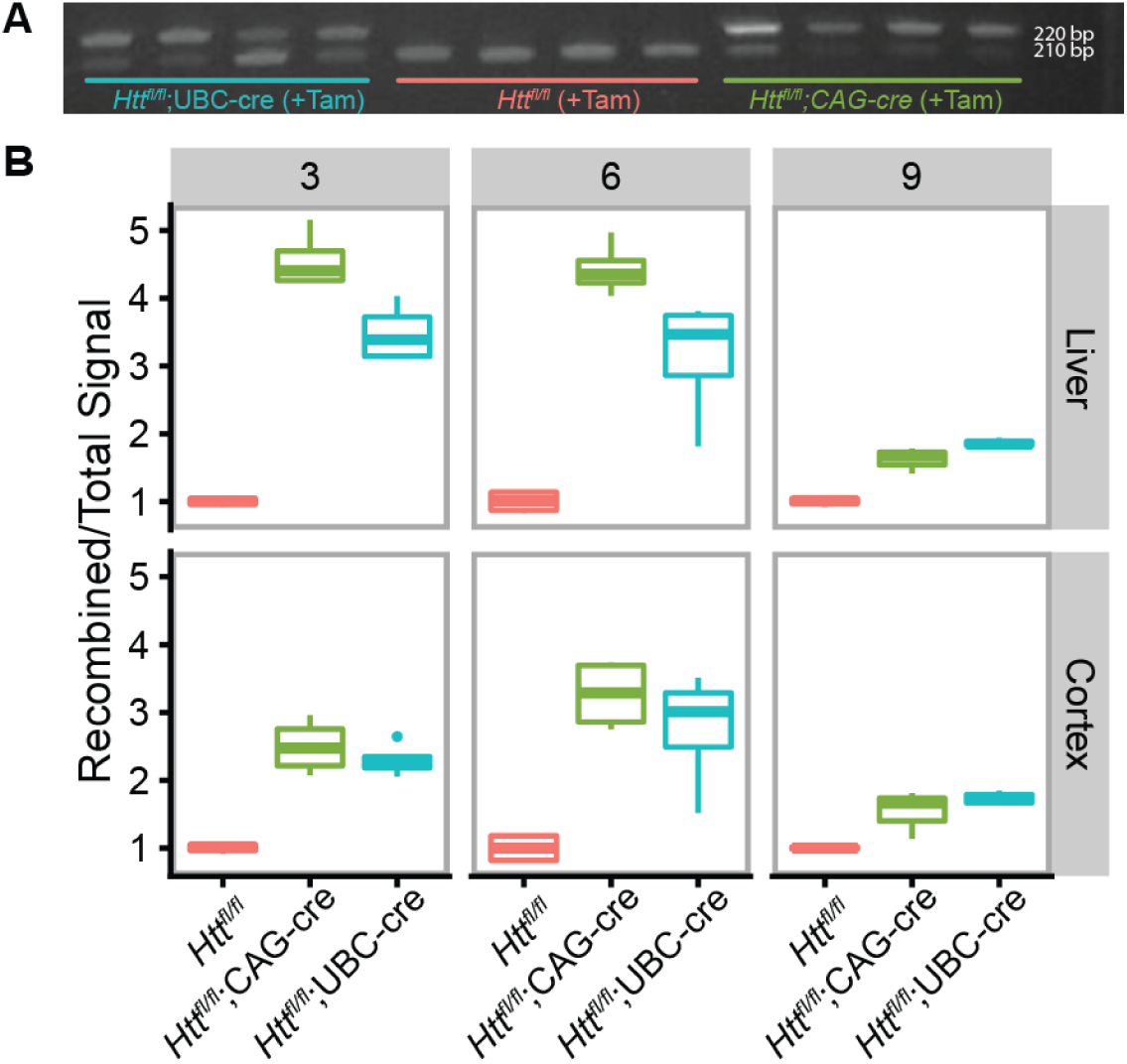
Semi-quantitative PCR demonstrates successful recombination of the *Htt* gene following tamoxifen administration. **A)** Example image of a gel used for quantification. The top band (220bp) is the recombined PCR product, visible only in tamoxifen-treated *Htt^fl/fl^*;CAG-cre and *Htt^fl/fl^*;UBC-cre animals, and the bottom band (210bp) is the unrecombined product, visible in all animals but fainter in lanes with more recombination. **B)** Recombination efficiency was calculated by dividing the intensity of the recombined band by the total signal of the recombined plus unrecombined bands. A higher value is equivalent to more recombination.

**Supplemental figure S6:**
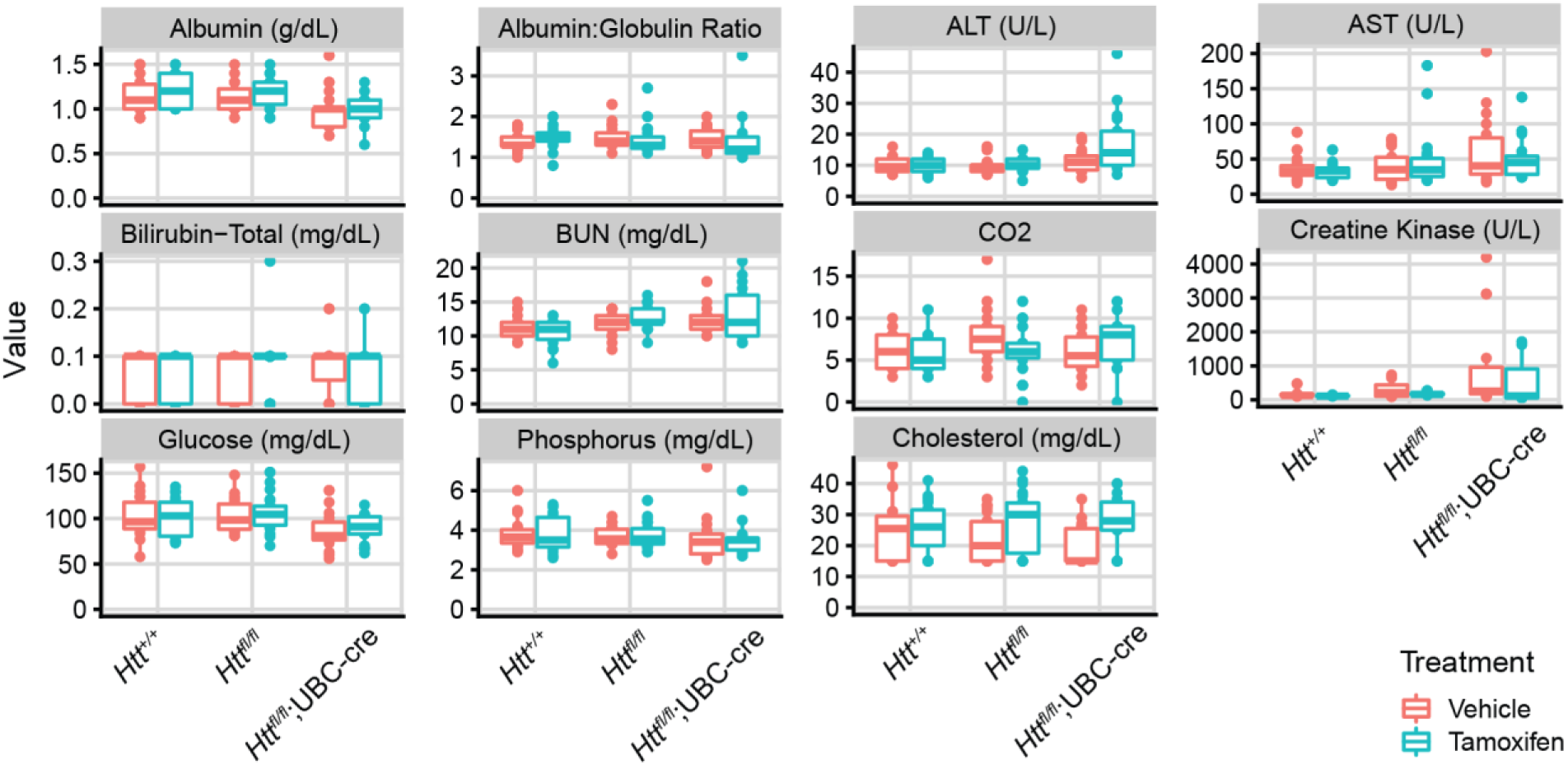
Plasma chemistry. Of the 11 plasma analytes measured, only alanine transaminase (ALT) showed any significant differences in response to Htt loss.

**Table S1.**
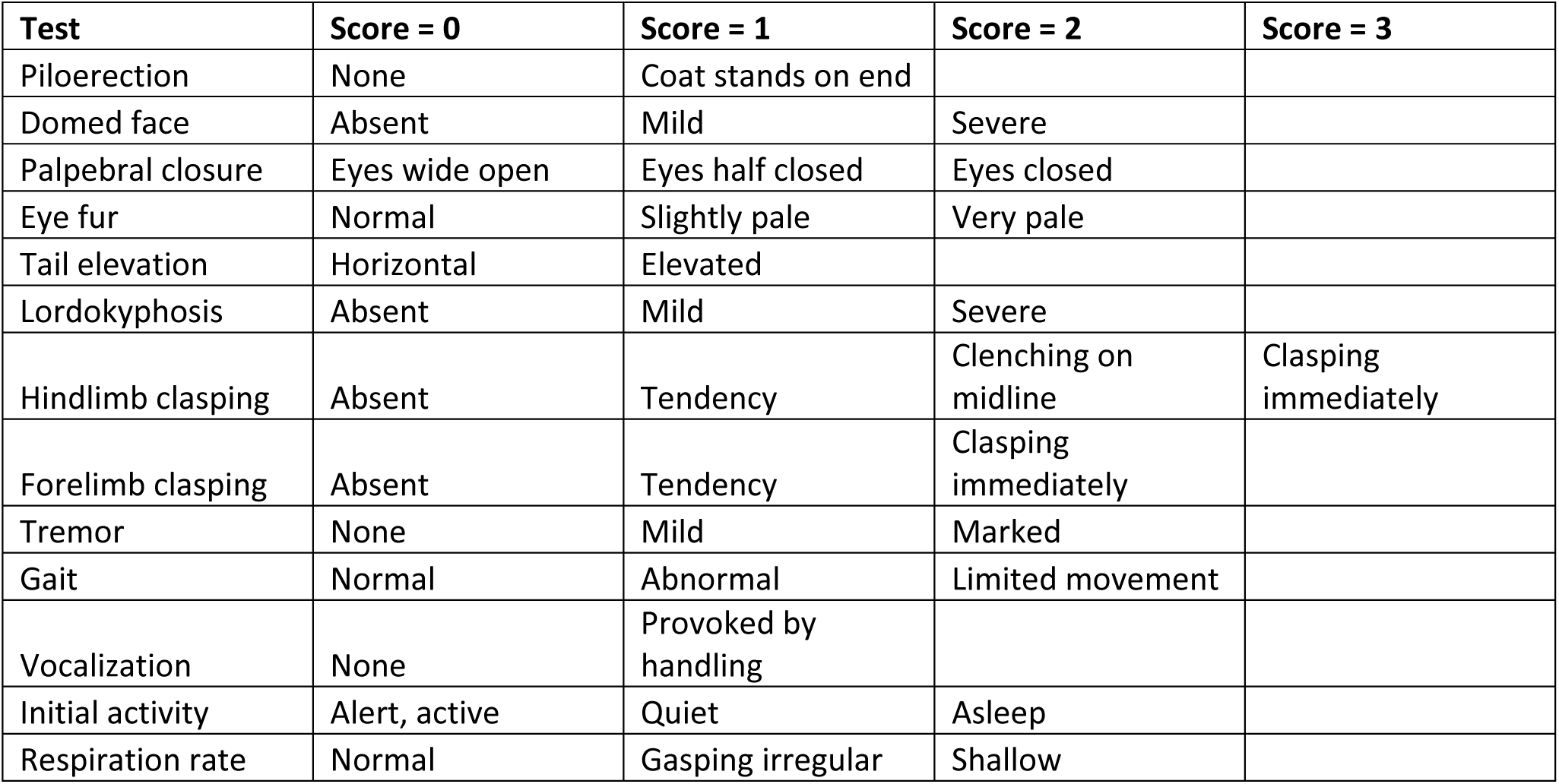
SHIRPA rubric.

## References

1. Bates GP, Dorsey R, Gusella JF, Hayden MR, Kay C, Leavitt BR, et al. Huntington disease. Nature reviews Disease primers. 2015;1:15005.

2. Duyao M, Auerbach A, Ryan A, Persichetti F, Barnes G, McNeil S, et al. Inactivation of the mouse Huntington’s disease gene homolog Hdh. Science. 1995;269(5222):407–10.

3. Nasir J, Floresco SB, O’Kusky JR, Diewert VM, Richman JM, Zeisler J, et al. Targeted disruption of the Huntington’s disease gene results in embryonic lethality and behavioral and morphological changes in heterozygotes. Cell. 1995;81(5):811–23.

4. Zeitlin S, Liu JP, Chapman DL, Papaioannou VE, Efstratiadis A. Increased apoptosis and early embryonic lethality in mice nullizygous for the Huntington’s disease gene homologue. Nat Genet. 1995;11(2):155–63.

5. Rodan LH, Cohen J, Fatemi A, Gillis T, Lucente D, Gusella J, et al. A novel neurodevelopmental disorder associated with compound heterozygous variants in the huntingtin gene. Eur J Hum Genet. 2016;24(12):1826–7.

6. Lopes F, Barbosa M, Ameur A, Soares G, Sá J de, Dias AI, et al. Identification of novel genetic causes of Rett syndrome-like phenotypes. Journal of medical genetics. 2016;53(3):190–9.

7. Karczewski KJ, Francioli LC, Tiao G, Cummings BB, Alföldi J, Wang Q, et al. The mutational constraint spectrum quantified from variation in 141,456 humans. Nature. 2020;581(7809):434–43.

8. Tabrizi SJ, Ghosh R, Leavitt BR. Huntingtin Lowering Strategies for Disease Modification in Huntington’s Disease. Neuron. 2019;101(5):801–19.

9. Keller CG, Shin Y, Monteys AM, Renaud N, Beibel M, Teider N, et al. An orally available, brain penetrant, small molecule lowers huntingtin levels by enhancing pseudoexon inclusion. Nat Commun. 2022;13(1):1150.

10. Bhattacharyya A, Trotta CR, Narasimhan J, Wiedinger KJ, Li W, Effenberger KA, et al. Small molecule splicing modifiers with systemic HTT-lowering activity. Nat Commun. 2021;12(1):7299.

11. Claassen DO, Corey-Bloom J, Dorsey ER, Edmondson M, Kostyk SK, LeDoux MS, et al. Genotyping single nucleotide polymorphisms for allele-selective therapy in Huntington disease. Neurology Genetics. 2020;6(3):e430.

12. Tabrizi SJ, Leavitt BR, Landwehrmeyer GB, Wild EJ, Saft C, Barker RA, et al. Targeting Huntingtin Expression in Patients with Huntington’s Disease. New Engl J Med. 2019;380(24):2307–16.

13. Raamsdonk JMV, Gibson WT, Pearson J, Murphy Z, Lu G, Leavitt BR, et al. Body weight is modulated by levels of full-length Huntingtin. Hum Mol Genet. 2006;15(9):1513–23.

14. Wang G, Liu X, Gaertig MA, Li S, Li XJ. Ablation of huntingtin in adult neurons is nondeleterious but its depletion in young mice causes acute pancreatitis. Proc National Acad Sci. 2016;113(12):3359– 64.

15. Dietrich P, Johnson IM, Alli S, Dragatsis I. Elimination of huntingtin in the adult mouse leads to progressive behavioral deficits, bilateral thalamic calcification, and altered brain iron homeostasis. Plos Genet. 2017;13(7):e1006846.

16. Barro C, Chitnis T, Weiner HL. Blood neurofilament light: a critical review of its application to neurologic disease. Ann Clin Transl Neur. 2020;7(12):2508–23.

17. Hayashi S, McMahon AP. Efficient Recombination in Diverse Tissues by a Tamoxifen-Inducible Form of Cre: A Tool for Temporally Regulated Gene Activation/Inactivation in the Mouse. Dev Biol. 2002;244(2):305–18.

18. Quintana A, Kruse SE, Kapur RP, Sanz E, Palmiter RD. Complex I deficiency due to loss of Ndufs4 in the brain results in progressive encephalopathy resembling Leigh syndrome. Proc Natl Acad Sci. 2010;107(24):10996–1001.

19. Ruzankina Y, Pinzon-Guzman C, Asare A, Ong T, Pontano L, Cotsarelis G, et al. Deletion of the Developmentally Essential Gene ATR in Adult Mice Leads to Age-Related Phenotypes and Stem Cell Loss. Cell Stem Cell. 2007;1(1):113–26.

20. Heffner C. INTRAPERITONEAL INJECTION OF TAMOXIFEN FOR INDUCIBLE CRE-DRIVER LINES [Internet]. 2011 [cited 2018 Nov 1]. Available from: https://www.jax.org/research-and-faculty/resources/cre-repository/tamoxifen

21. Bragg RM, Coffey SR, Cantle JP, Hu S, Singh S, Legg SR, et al. Huntingtin loss in hepatocytes is associated with altered metabolism, adhesion, and liver zonation. Life Sci Alliance. 2023;6(11):e202302098.

22. Science AI for B. Allen Mouse Brain Atlas [dataset] [Internet]. 2011. Available from: mouse.brain-map.org

23. Tran D, DiGiacomo P, Born DE, Georgiadis M, Zeineh M. Iron and Alzheimer’s Disease: From Pathology to Imaging. Front Hum Neurosci. 2022;16:838692.

24. Nakamura T, Naguro I, Ichijo H. Iron homeostasis and iron-regulated ROS in cell death, senescence and human diseases. Biochim Biophys Acta (BBA) - Gen Subj. 2019;1863(9):1398–409.

25. Liu JP, Zeitlin SO. Is Huntingtin Dispensable in the Adult Brain? J Huntington’s Dis. 2017;Preprint(Preprint):1–17.

26. Kaemmerer WF, Grondin RC. The effects of huntingtin-lowering: what do we know so far? Degener Neurological Neuromuscul Dis. 2019;9:3–17.

27. Auerbach W, Hurlbert MS, Hilditch-Maguire P, Wadghiri YZ, Wheeler VC, Cohen SI, et al. The HD mutation causes progressive lethal neurological disease in mice expressing reduced levels of huntingtin. Hum Mol Genet. 2001;10(22):2515–23.

28. Boudreau RL, McBride JL, Martins I, Shen S, Xing Y, Carter BJ, et al. Nonallele-specific Silencing of Mutant and Wild-type Huntingtin Demonstrates Therapeutic Efficacy in Huntington’s Disease Mice. Mol Ther. 2009;17(6):1053–63.

29. Spronck EA, Vallès A, Lampen MH, Montenegro-Miranda PS, Keskin S, Heijink L, et al. Intrastriatal Administration of AAV5-miHTT in Non-Human Primates and Rats Is Well Tolerated and Results in miHTT Transgene Expression in Key Areas of Huntington Disease Pathology. Brain Sci. 2021;11(2):129.

30. McBride JL, Pitzer MR, Boudreau RL, Dufour B, Hobbs T, Ojeda SR, et al. Preclinical Safety of RNAi-Mediated HTT Suppression in the Rhesus Macaque as a Potential Therapy for Huntington’s Disease. Mol Ther. 2011;19(12):2152–62.

31. Grondin R, Kaytor MD, Ai Y, Nelson PT, Thakker DR, Heisel J, et al. Six-month partial suppression of Huntingtin is well tolerated in the adult rhesus striatum. Brain. 2012;135(4):1197–209.

32. Stiles DK, Zhang Z, Ge P, Nelson B, Grondin R, Ai Y, et al. Widespread suppression of huntingtin with convection-enhanced delivery of siRNA. Experimental neurology. 2012;233(1):463–71.

33. Grondin R, Ge P, Chen Q, Sutherland JE, Zhang Z, Gash DM, et al. Onset Time and Durability of Huntingtin Suppression in Rhesus Putamen After Direct Infusion of Antihuntingtin siRNA. Mol Ther - Nucleic Acids. 2015;4(6):e245.

34. Yang S, Chang R, Yang H, Zhao T, Hong Y, Kong HE, et al. CRISPR/Cas9-mediated gene editing ameliorates neurotoxicity in mouse model of Huntington’s disease. J Clin Invest. 2017;

35. Shin JW, Hong EP, Park SS, Choi DE, Zeng S, Chen RZ, et al. PAM-altering SNP-based allele-specific CRISPR-Cas9 therapeutic strategies for Huntington’s disease. Mol Ther - Methods Clin Dev. 2022;26:547–61.

36. Monteys AM, Ebanks SA, Keiser MS, Davidson BL. CRISPR/Cas9 Editing of the Mutant Huntingtin Allele In Vitro and In Vivo. Mol Ther. 2017;25(1):12–23.

37. Kolli N, Lu M, Maiti P, Rossignol J, Dunbar GL. CRISPR-Cas9 Mediated Gene-Silencing of the Mutant Huntingtin Gene in an In Vitro Model of Huntington’s Disease. Int J Mol Sci. 2017;18(4):754.

38. Mortberg MA, Gentile JE, Nadaf NM, Vanderburg C, Simmons S, Dubinsky D, et al. A single-cell map of antisense oligonucleotide activity in the brain. Nucleic Acids Res. 2023;51(14):7109–24.

39. Dragatsis I, Levine MS, Zeitlin S. Inactivation of Hdh in the brain and testis results in progressive neurodegeneration and sterility in mice. Nat Genet. 2000;26(3):ng1100_300.

40. Dietrich P, Shanmugasundaram R, E S, Dragatsis I. Congenital hydrocephalus associated with abnormal subcommissural organ in mice lacking huntingtin in Wnt1 cell lineages. Hum Mol Genet. 2009;18(1):142–50.

41. Yang G, Nowsheen S, Aziz K, Georgakilas AG. Toxicity and adverse effects of Tamoxifen and other anti-estrogen drugs. Pharmacol Ther. 2013;139(3):392–404.

42. Crow YJ, Stetson DB. The type I interferonopathies: 10 years on. Nat Rev Immunol. 2021;1–13.

43. Crow YJ, Manel N. Aicardi–Goutières syndrome and the type I interferonopathies. Nat Rev Immunol. 2015;15(7):429–40.

44. Koob M, Laugel V, Durand M, Fothergill H, Dalloz C, Sauvanaud F, et al. Neuroimaging In Cockayne Syndrome. Am J Neuroradiol. 2010;31(9):1623–30.

45. Ieshima A, Kisa T, Yoshino K, Takashima S, Takeshita K. A morphometric CT study of Down’s syndrome showing small posterior fossa and calcification of basal ganglia. Neuroradiology. 1984;26(6):493–8.

46. Wisniewski KE, French JH, Rosen JF, Kozlowski PB, Tenner M, Wisniewski Hm. Basal Ganglia Calcification (BGC) in Down’s Syndrome (DS)—Another Manifestation of Premature Aging*. Ann Ny Acad Sci. 1982;396(1):179–89.

47. Peters MEM, Brouwer EJM de, Bartstra JW, Mali WPTM, Koek HL, Rozemuller AJM, et al. Mechanisms of calcification in Fahr disease and exposure of potential therapeutic targets. Neurology Clin Pract. 2019;10(5):449–57.

48. Wang C, Li Y, Shi L, Ren J, Patti M, Wang T, et al. Mutations in SLC20A2 link familial idiopathic basal ganglia calcification with phosphate homeostasis. Nat Genet. 2012;44(3):254–6.

49. Legati A, Giovannini D, Nicolas G, López-Sánchez U, Quintáns B, Oliveira JRM, et al. Mutations in XPR1 cause primary familial brain calcification associated with altered phosphate export. Nat Genet. 2015;47(6):579–81.

50. Keller A, Westenberger A, Sobrido MJ, García-Murias M, Domingo A, Sears RL, et al. Mutations in the gene encoding PDGF-B cause brain calcifications in humans and mice. Nat Genet. 2013;45(9):1077–82.

51. Nicolas G, Pottier C, Maltête D, Coutant S, Rovelet-Lecrux A, Legallic S, et al. Mutation of the PDGFRB gene as a cause of idiopathic basal ganglia calcification. Neurology. 2013;80(2):181–7.

52. Dietrich P, Johnson IM, Alli S, Dragatsis I. Elimination of huntingtin in the adult mouse leads to progressive behavioral deficits, bilateral thalamic calcification, and altered brain iron homeostasis. PLoS Genet. 2017;13(7):e1006846.

53. Luong TN, Carlisle HJ, Southwell A, Patterson PH. Assessment of Motor Balance and Coordination in Mice using the Balance Beam. J Vis Exp. 2011;(49).

54. Fischer DF, Dijkstra S, Lo K, Suijker J, Correia ACP, Naud P, et al. Development of mAb-based polyglutamine-dependent and polyglutamine length-independent huntingtin quantification assays with cross-site validation. PLoS ONE. 2022;17(4):e0266812.

